# Two decades of Novel Ecosystems: Literature analysis of topics, trends and connectivity

**DOI:** 10.64898/2026.10.09.758140

**Authors:** Timothy L. Staples, Peter Bridgewater, Yvonne Buckley, Giselda Durigan, Jim Harris, Tina Heger, Matthew Kerr, Sybryn L. Maes, Siân Moxon, Stephen Murphy, Alejandro Ordonez, Rachel Standish, Jens-Christian Svenning, Brenda Maria Zoderer, Marcus Collier

**Affiliations:** School of the Environment, The University of Queensland, Australia; Centre for Conservation Ecology and Genomics, University of Canberra, Australia; School of Natural Sciences, Trinity College Dublin; Instituto de Pesquisas Ambientais, SP, Brazil; BOKU University, Department of Landscape, Water and Infrastructure, Institute of Landscape Development, Recreation and Conservation Planning, Vienna, Austria; Cranfield Environment Centre, Cranfield University, Bedfordshire, MK430AL, UK; Technical University of Munich, Germany; TUM School of Life Sciences, Restoration Ecology, Emil-Ramann-Str. 6, 85354 Freising, Germany; Leuphana University Lüneburg, Institute of Ecology, Universitätsallee 1, 21335 Lüneburg, Germany; Center for Ecological Dynamics in a Novel Biosphere (ECONOVO), Department of Biology, Aarhus University, Aarhus C, Denmark; Division of Forest, Nature & Landscape, KU Leuven, Leuven, Belgium; School of Art, Architecture and Design, London Metropolitan University, London, UK; School of Environment, Resources & Sustainability, University of Waterloo, Waterloo ON Canada; School of Environmental and Conservation Sciences, Murdoch University, Perth, Western Australia

**Keywords:** Bibliometrics, ecosystem management, ecological novelty, ecological restoration, linguistic analysis, topic modelling

## Abstract

1. As human-driven climate change and other anthropogenic forces produce widespread impacts on biodiversity and ecosystem functioning, many ecosystems are shifting into unprecedented, novel states. The Novel Ecosystem Concept (NEC), formalized in 2006, was developed as an alternative framework for managing “novel ecosystems”, human-altered systems that are challenging to restore to their historical state. In the years since, the NEC has been proposed as a broader lens for ecosystem management under ongoing global change.
2. We evaluated how the uptake and potential of the NEC have been realized over time using a bibliometric and linguistic toolkit. Using bibliometric data from the Web of Science, we clustered 40,914 applied ecology papers published between 2006 and 2025 into twenty research topics based on language use. We quantified the volume of NEC research within each topic, rates of publication over time, and cross-topic connectivity proxied by citations.
3. We found NEC research in most of the topics of applied ecology, particularly invasion biology, ecosystem services, soil research and studies of freshwater, urban and marine systems. In contrast, several major conservation-focused topics, including protected areas, threatened species, and human–wildlife interactions, contained little research with an explicit link to novel ecosystems. Publication rates increased across all topics, although growth slowed over time. At the same time, NEC research became more focused into ecosystem-specific topics, especially marine ecosystems. NEC research across different topics was well-connected by citations, with papers more likely to cite each other when they used similar language.
4. Overall, the NEC does not appear to have expanded beyond its origins in restoration to become a broadly applied framework across conservation and ecosystem management. Instead, the concepts behind the NEC have been integrated into restoration and management frameworks grappling with climate and other anthropogenic change, while “novelty” is emerging as a powerful quantitative variable for measuring ecological change over space and time. Our results highlight the potential value of bibliometric and linguistic tools to deconstruct the ecological literature to inform what we can, or should, do to intervene for benefit of nature and people.

---

There has never been a greater need for science to inform ecosystem intervention and management. Biosphere-scale crises are looming: climate change and human landscape modification are threatening fundamental ecosystem integrity and driving species extinctions (Jaureguiberry et al., 2026). In response, the United Nations has outlined key biodiversity targets (Secretariat of the Convention on Biological Diversity, 2022) and declared consecutive decades on biodiversity conservation (United Nations General Assembly, 2010) and ecosystem restoration (United Nations General Assembly, 2019). Conservation is a proactive process that aims to mitigate anthropogenic processes that could alter or degrade ecosystems (Sandbrook, 2015). Restoration complements conservation, intervening in degraded systems to reverse the loss of biodiversity and recover ecological structure and function (Jones et al., 2018). As climate change and other human pressures amplify, both these endeavours are becoming more challenging (Harris et al., 2006; Lawler, 2009; Maes et al., 2024). Climate change is impacting all ecosystems, regardless of conservation protection (Araújo et al., 2004) and restoring some degraded systems back to their historic state can be challenging and, in some cases, infeasible (Brudvig, 2017; Jackson & Hobbs, 2009). The emergence of novel ecosystems and climates create challenges for conventional management frameworks that prioritize maintenance and recovery of pre-human ecosystem states (Lindenmayer et al., 2008; Williams & Jackson, 2007). Successful conservation and restoration hinges on updating and adapting how we consider ecosystem management in light of intensifying human impacts (Hobbs et al., 2011; Williams et al., 2021).

The Novel Ecosystem Concept (NEC) was first introduced following a UNESCO/SCOPE workshop on emerging ecosystems held in 2000, designed to improve management outcomes in an increasingly human-altered world (Hobbs et al., 2006). While both conservation and restoration have relied on the use of historical reference states to orient management (McNellie et al., 2020), the NEC argues that past ecosystem states are sometimes undesirable or unachievable management goals (Hobbs et al., 2014), built on three key components. First, some unmanaged ecosystems are in a “novel” state, with species and abiotic regimes that do not resemble past or nearby reference systems. Examples of novel ecosystems include post-mine sites (Jagodziński et al., 2018), drained wetlands (Kreyling et al., 2021) and invaded islands (Mascaro et al., 2012). Second, these novel ecosystems have been co-produced by both natural and anthropogenic forces. Third, in some cases avoiding, or reversing, human impacts in these systems may not be possible, feasible or even desirable (Hobbs et al., 2013; Williams & Brown, 2024). These barriers to restoration success can be ecological, with human-altered states locked in due to regime shifts, hysteresis and positive feedback loops (Cipriotti et al., 2019; Maes et al., 2024; Österblom et al., 2007; Sha et al., 2026). Restoring the historic ecosystem state can also be technically achievable, but infeasible within social or economic limits (Mañas-Navarro et al., 2023; zu Ermgassen & Löfqvist, 2024). The NEC provides an alternative lens for ecological restoration that directs management goal-setting to maximize success and ecological return on investment (Evers et al., 2018; Hallett et al., 2013), often by focusing on restoring important ecosystem functions or species interactions instead of holistic reconstruction of the reference ecosystem (Hulvey et al., 2013). Given this reframing, the NEC has potential to provide an ecosystem-agnostic perspective to approach management in a human-dominated future (Hobbs et al., 2009).

In the twenty years since initial publication in 2006, the reception of the NEC has been mixed. By classic metrics of scientific success, seminal Novel Ecosystem papers (Hobbs et al., 2006, 2009, 2014) have been well-cited and attracted follow-on funding and publications (Hobbs, 2017). There is growing interest in the impacts of climate change on management (Simonson et al., 2021), and on measuring and understanding “ecological novelty” (Kerr et al., 2026; Staples et al., 2025). Despite this interest, conservation biologists critiqued a vague and inoperable definition of “novel” that “lowered the bar” for bad ecological actors to avoid restoration responsibilities (Aronson et al., 2014; Murcia et al., 2014). These critiques may reflect broader research segregation, as conservation, restoration and climate change research largely exist within parallel and non-overlapping bodies of the ecological literature (Staples et al., 2019; Young, 2000). The NEC has broad potential to help manage a planet undergoing widespread environmental change (Hobbs, 2017; Radeloff et al., 2015) but it remains unclear whether, and how, the NEC has informed, and been integrated into, applied ecological research.

Bibliometrics, the meta-study of patterns in the academic literature, may be a useful diagnostic tool for understanding how, and how broadly, the potential of the NEC has been realised. Text-based clustering models (“topic models”) can decompose complex research landscapes into core topics based on the language use within papers (e.g., Zhang et al., 2017). Estimating NEC publication trends across the major topics of applied ecology can provide an overall measure of impact and influence, including identifying research areas with little NEC engagement. In addition to publications, citations can serve as a proxy of research interest (Bornmann & Daniel, 2008). In well-connected research, the number of citations between a pair of topics should be predictable based on two factors, the size of each topic and the linguistic similarity between the two topics, on the assumption that language similarity translates into research overlap (Dias et al., 2018). Leftover variation in citations after accounting for these hypothetical factors can highlight topics that are overly well-connected, and conversely poorly connected, to the broader literature. Together, bibliometric and linguistic tools may provide a powerful way to chart the reach and impact of the NEC since 2006 and identify how much of its potential value as a lens for ecosystem management has been been realized.

Here we interrogated the applied ecology literature to ask, in the words of one of the original authors of the NEC, whether it has been “a new direction, or a road to nowhere” (Hobbs, 2017). With bibliometric data obtained from the Web of Science™ Core Collection published by Clarivate, we asked:

1. What are the major topics in applied ecological research, and how is NEC research distributed across these topics?
2. How has research on the NEC changed over time across major topics, in both publication rates and relative topic rankings?
3. How well can citations between NEC research be predicted by topic size and linguistic similarity?, and
4. Is variation in citation counts (after size-and-similarity adjustment) conserved within topics, providing an indication of overall topic connectivity to the broader NEC literature?

We hypothesized that (1) research on the NEC would be skewed towards topics on direct management and intervention in ecosystems, reflecting the focus of initial publications (Hobbs et al., 2009). We anticipated that conservation topics would contain little research on the NEC, perpetuating early criticisms (Aronson et al., 2014; Murcia et al., 2014). Given our study period of only nineteen years (time since initial NEC publication), we expected (2) rates of NEC publication to remain relatively stable across topics, assuming that topics that found the NEC valuable would likely continue study over time. Finally, we hypothesized (3) that research on the NEC would be well-connected across topics. We expected the number of citations between topic pairs to be high overall, increasing with more similar language use between topics and with larger topic size, both in the number of NEC papers and size in broader applied ecology. We expected that (4) leftover variation in citation counts, after accounting for topic size and similarity correlations, would be conserved within topics and act as a useful indicator of whether topics were well-connected and poorly connected to the broader NEC literature.

## Methods

Our methodology involved identifying the NEC literature, which we accessed via the Web of Science™ (WoS) Core Collection published by Clarivate™. We then used the titles, keywords and abstracts of papers in a linguistic topic association model to identify core topics. These topics were used to address our research questions. First, we modelled the abundance of NEC papers, and the papers citing them, within each topic. Second, we modelled the rate of new publications in each topic across our study period. And finally, we modelled how well we could predict citation counts between topics based on their size and similarity of language.

### Publication and citation collection

We downloaded publication data from WoS on 27th February 2026 (Fig. 1).

**Figure 1:**
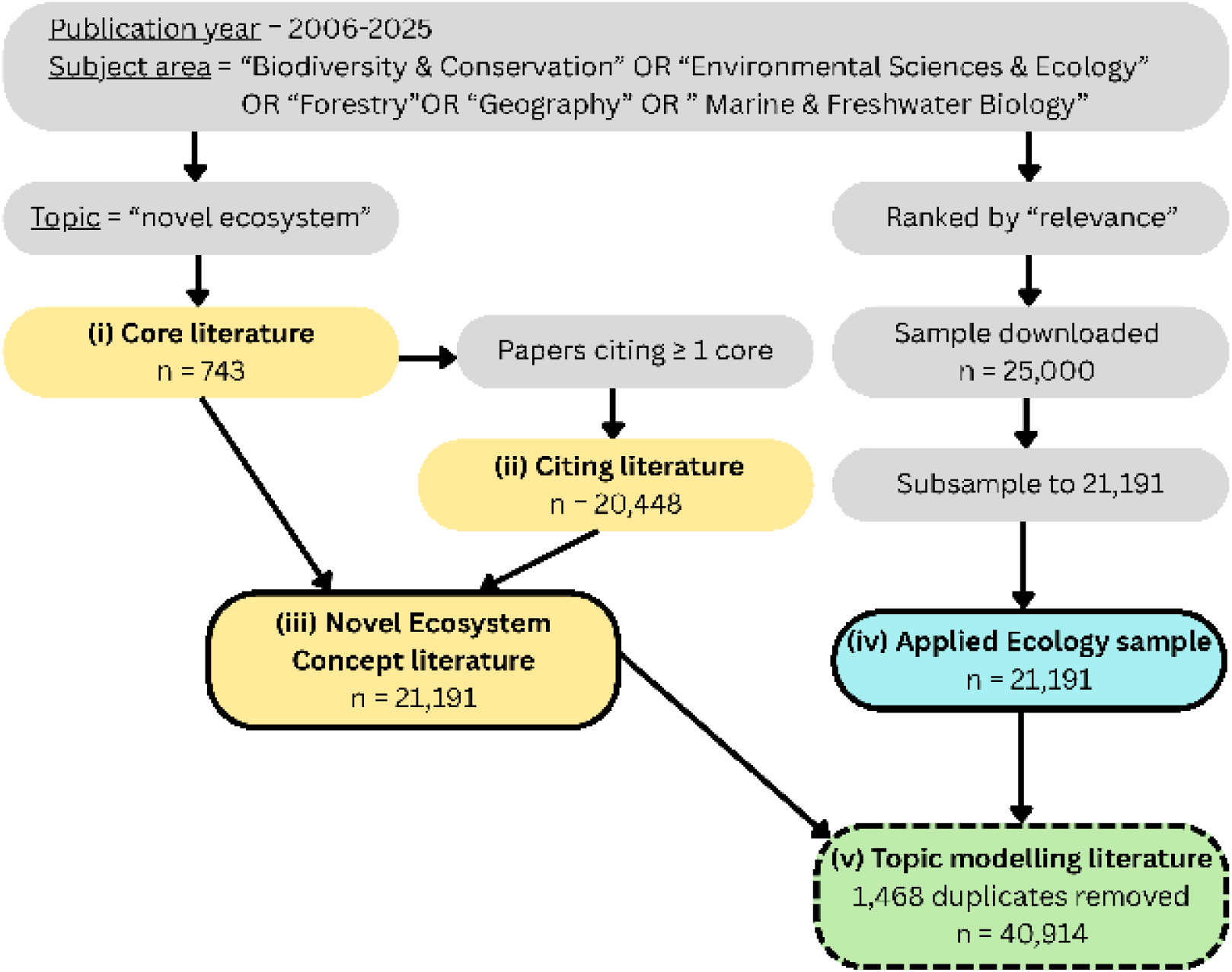
Outline of methodology used to access Web of Science publication data. Numerals indicate literature groups referred to in analyses. The “Topic modelling literature” (dashed outline) is an aggregate group only used for the topic model to cluster papers based on language use.

First, we downloaded publications to represented the NEC literature. This dataset comprised two sets of papers. The first included 743 “core papers” that mentioned novel ecosystems in their title, abstract or keywords, published between 2006 and 2025 and within a set of ecological subject areas (i in Fig. 1). To capture broader interest in NEC research, we added an additional 20,448 “citing papers” that cited at least one core paper (ii in Fig. 1). We refer to the combined core and citing papers (n = 21,191) as the “Novel Ecosystem Concept literature” (NEC literature: iii in Fig. 1).

Using just the NEC literature may have caused our topic modelling approach to miss research topics in applied ecology that were not represented in the NEC literature. To account for this, we downloaded a subset of broader applied ecology research from WoS to act as a control group, sized to 21,191 papers to match the NEC literature (iv in Fig. 1). Our goal was not to exhaustively include every paper in applied ecology, but rather to ensure our topic modelling did not exclude important sections of the literature that were “dark spots” for NEC research. We used the applied ecology sample three times (described in detail below): first, in the topic association modelling, combined with the NEC literature (v in Fig. 1), second, to contrast overall topic size (as a proportion of the sample) with volume of NEC research in each topic, and third, as a predictor variable in our topic citation model.

### Data analysis

All data processing and analyses were performed in R version 4.4.0 (R Core Team, 2024), using the following packages: betareg (Cribari-Neto & Zeileis, 2010), DHARMa (Hartig, 2020), glmmTMB (Brooks et al., 2017), lme4 (Bates et al., 2015), MASS (Venables & Ripley, 2002), tm (Feinerer et al., 2008), topicmodels (Grün & Hornik, 2011) and vegan (Oksanen et al., 2025). Assumptions of all statistical models were validated via residual simulation using the DHARMa package.

#### Topic Association Modelling

We conducted the topic association modelling on a linguistic corpus, combining the titles, keywords and abstracts of each paper into a single “text”, across both the NEC and applied ecology sample literature (Fig. 1). These are intentional fields where authors are incentivised to distil the clearest description of their paper’s content, making them ideal input data for a topic association model. We ran the topic model on the combined NEC literature and sample literature (n = 40,914), removing 1,468 duplicates where NEC papers were also in the sample literature (v in Fig. 1).

Prior to modelling, we cleaned each text by removing: (1) all punctuation and numerals, and converted all words to lowercase, (2) stopwords, small words with no semantic value (e.g., the, to, and) using the stopwords function in the tm package, (3) word suffices using a process called “stemming”, using the stemDocument function of the tm package (e.g., “ecology” and “ecological” both become “ecolog”), (4) publisher words that were sometimes appended to the end of the abstract field (e.g., “author”, “copyright”), (5) non-English language abstracts for dual-abstract papers and (6) words that were only used once across the entire corpus (n = 27.029). In total the dataset contained 7,278,872 samples of 42,382 unique word stems, with a median of 174 words per text.

The topic model we employed was a Latent Dirichlet Allocation (LDA). This model uses a Bayesian network to associate words, and texts, into a set number of topics (Jelodar et al., 2019; for an inuitive explanation, see Ganegedara, 2018). Words that commonly co-occur are linked into the same topic, which the model uses to estimate the probability each text falls into a given topic, based on the words it uses. The LDA is a soft-clustering algorithm, in that each paper was given a set of probabilities of falling into each identified topic, rather than simply a topic classification. We tested LDA models containing 5-30 topics, in units of 5, with 20 topics providing the best trade-off of complexity to explanatory power (Fig. S1).

We assigned each paper to a topic based on the highest probability within the LDA, retaining the topic-paper probability matrix to calculate topic language similarity. We named each topic by extracting the 15 most influential words that defined each topic (from the LDA) and papers with the highest likelihood of association with the topic, using our understanding of the literature augmented by a Large Language Model (Anthropic, 2026: Supplementary Methods).

Where topics reflected a broad research topic without a clear identity, we performed subsequent LDAs to identify subtopics. LDAs were fit as per above, using only papers within a topic and limited to 15 subtopics.

#### NEC distribution across the topics of applied ecology

To establish how research on the NEC was distributed across topics, we visually compared the number of core papers published within each topic to the proportional size of each topic in the applied ecology sample literature. We then used a negative binomial generalised linear model (glm.nb function, MASS package) to estimate the relationship between the number of core NEC papers and the number of citing papers. The response variable of this model was the count of citing papers for each topic. The sole fixed effect was natural log-transformed count of core papers in each topic.

#### Change in NEC research over time

We established topic growth rates over time using a negative binomial-distributed generalized linear model (glm.nb function, MASS package). This model used the count of core papers published in each topic in each year as the response variable. The model contained three fixed effects: publication year, year^2^ (to test for quadratic non-linear trends) and topic as a categorical fixed effect, interacting with both year terms. Predicted publication counts each year (from 2010 to 2025) were used to assign topic ranks to examine relative contribution of each topic to overall NEC research.

#### Citation connectivity between topics of the NEC

The topic association model detected topics solely by linguistic similarity, not by the strength of research connectivity as proxied by citations. We used cited reference information in the WoS data to calculate citation counts between every combination of “citing topic” and “cited topic”. excluding intra-topic pairs (n = 380). We modelled these citation counts to identify which topic pairs were well, or poorly connected by citations, based on their linguistic similarity, as well as each topic’s size within the sample literature and the NEC literature.

We obtained citation counts by matching DOI numbers in the “cited references” field of WoS data for each paper in the NEC literature. We identified a total of 1,816,234 citations made by the 21,191 papers. Of these, 128,690 citations (7.09%) were made to other NEC papers. We summed NEC citations from each topic (“citing topic”) to each topic (“cited topic”), excluding intra-topic citation counts, yielding 380 topic-topic citation counts.

These citation counts were used as the response variable in two negative binomial generalized mixed-effects models (glmmTMB function, nbinom2 distribution, glmmTMB package).

The first model was a “size-correction model”, fit with four fixed effects to correlate citation counts with the size of cited and citing topics. The first two fixed effects were size of the cited topic and citing topic in the sample literature, fit as a proportion of the sample literature size of 21,191, natural log-transformed and fit with an interaction effect. The other two fixed effects were the number of core NEC papers in the cited topic and citing topic, also natural log-transformed with an interaction effect.

The second model, a “linguistic similarity” model, added linguistic similarity as an additional fixed effect to the size-correction model. This two-model approach allowed us to compare models and estimate how well language similarity predicted citation counts after accounting for size-related sampling effects. We estimated language similarity using the gamma matrix from the LDA topic model, which contained a per-paper probability of being assigned to each topic, with the topic probabilities for a single paper summing to one (n = 40,914 papers x 20 topics). We used this matrix to calculate Bray-Curtis dissimilarity of paper topic probabilities between topics (vegdist function, vegan package), corrected to similarity by subtracting dissimilarities from one. Topic pairs with high similarities had correlated paper probabilities, an indication that language use between the two topics was more overlapping.

Both models were fit with two sets of crossed random intercepts, describing the (1) citing topic and (2) the cited topic of each topic pair. These random intercepts performed two functions. First, they reflected the nested nature of our citation data, given that each topic was the cited and citing topic 19 times each (one for each other topic excluding itself). This nesting violates the assumption of independence in regression models. Second, these random intercepts provided an indication of which topics generated more citations (the random intercepts for citing a topic) or which topics received more citations (the random intercepts for having cited a topic). The combination of these random intercepts provided a useful topic classification that we used to deconstruct topic connectivity in the NEC literature. Topics with high cited and citing random intercepts consistently made and received more citations to other NEC papers than predicted by the model, an indication they were central “hubs” of the literature. By contrast, topics with low cited and citing random intercepts consistently made and received few citations to other NEC papers and reflected “spokes” that were poorly connected to the broader literature. Topics with low citing random intercepts but high cited random intercepts were well-cited by NEC papers but did not cite the NEC literature in return. These topics acted as “authorities” that the NEC integrated into its research agenda without reciprocal citation from the authority topic. Finally, topics with high citing random intercepts but low cited random intercepts made citations to other NEC papers but were not cited in return. These topics represented “scouts” that were well-connected to the NEC literature but undercited by other topics.

## Results

### Topics of applied ecology

The twenty content topics detected within the applied ecology literature covered several major categories (Fig. 2, Table S1). Two topics, “Community & functional ecology” and “Ecological modelling & statistics”, were conceptual and taxa- and system-agnostic, aimed at understanding general ecological patterns. Five topics were defined by ecosystem (or part thereof) under study: “Forests & forestry”, “Freshwater & wetlands”, Marine & coastal ecology”, “Soil carbon & nutrients” and “Urban ecology”. Two topics were defined by taxa: “Wildlife ecology” and “Threatened species”. Five topics studied drivers of ecological change: “Fire & disturbance”, “Global change biology”, “Human-wildlife interactions”, “Invasion biology” and “Land use and landscapes”. Four topics were defined by the type of ecosystem management: “Conservation science”, “Ecological restoration”, “Ecosystem services” and “Protected areas”. The remaining two topics studied ecological systems through unique lenses. “Species distributions” examined patterns of species occurrence using species distribution and habitat modelling. “Regional studies” were country- or region-specific studies of ecological patterns for ecosystem management. Five of these topics exceeded 5% of the sample literature (point size and labels in Fig. 2): Conservation science, Human-wildlife interactions, Threatened species, Protected areas and Wildlife ecology. Only Conservation science exceeded 10% of the sample literature (Fig. 2), a meta-topic studying biodiversity and human relationships with conservation (Table S2).

**Figure 2:**
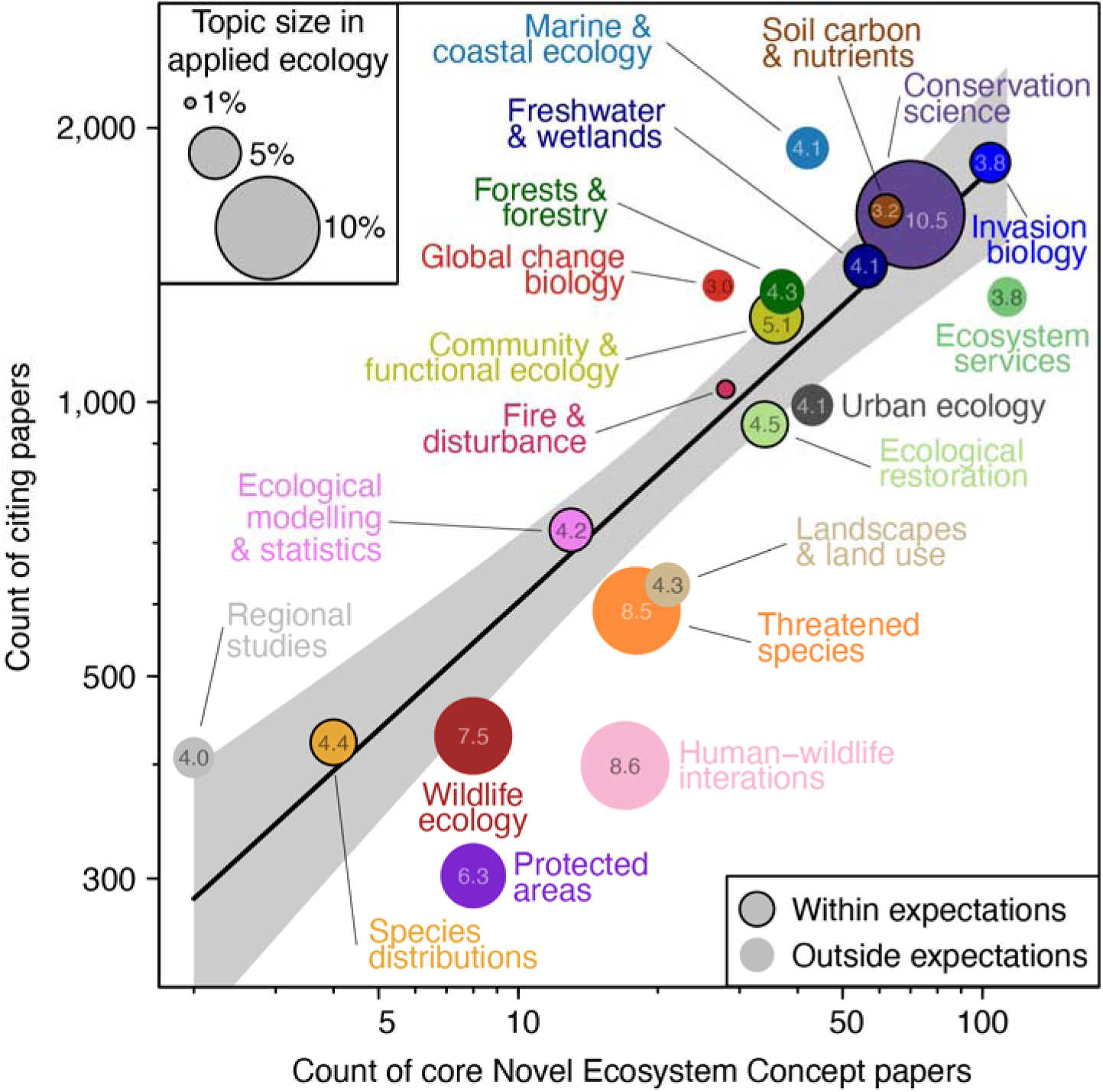
Applied ecology literature topics identified by Latent Dirichlet Allocation using paper titles, keywords and abstracts (Table S1), displayed as a function of the count of core Novel Ecosystem Concept papers (using the term “novel ecosystem”) published since 2006 (x-axis) and the count of papers that cited at least one core paper (y-axis). Point size and accompanying numbers reflect estimated topic size in broader applied ecology literature. The trend line represents the predicted citing paper count for a given core paper count from a negative binomial model, with the shaded region reflecting 95% confidence intervals (Table S3). Topics that fell below the lower 95% confidence interval had fewer citing papers than predicted from the model using their core paper count, and vice versa (“outside expectations”).

### NEC distribution across the topics of applied ecology

Five topics contained more than 50 core NEC publications (x-axis in Fig. 2): Ecosystem Services (n = 113), Invasion biology (n = 104), Conservation Science (n = 70), Soil carbon & nutrients (n = 62) and Freshwater & wetlands (n = 56). Core NEC papers in the Conservation Science topic were concentrated into research on Conservation in the Anthropocene (n = 18), Conservation ethics (n = 11) and Indigenous land management practices (n = 8: Table S2). NEC research in the Ecosystem Services topic was identified as a discrete sub-topic comprising 38 papers (5.1% of the 743 core papers) as well as Historical ecology (n = 19) and Resilience under climate change (n = 10: Table S3). Seven more topics, Community & functional ecology, Ecological restoration, Fire & disturbance, Forests & forestry, Global change biology, Marine & coastal ecology and Urban ecology contained more than 20 NEC papers (Fig. 2). The remaining seven topics, which included specific sub-disciplines of conservation research, all contained fewer than twenty core papers (Fig. 2).

Core paper count in a topic was a strong positive predictor of the number of citing papers; every 1% increase in core paper count corresponded with a 0.46% increase in citing paper count (Nagelkerke’s R^2^ = 0.951: Fig. 2, Table S4). Four topics had more citing papers than expected from their core paper count (exceeding the upper 95% confidence interval of the modelled trend). Forests & forestry and Regional studies were close to the upper confidence interval, while Global change biology and Marine & coastal ecology had 39% and 61% more citing papers than expected from the modelled trend, respectively (Fig. 2). Seven topics had fewer citing papers than expected from their core paper count (exceeding the lower 95% confidence interval of model trend: Fig. 2). Urban ecology and Wildlife ecology were close to the lower confidence interval, with citing paper counts within 83 and 79% of model predictions, respectively (Fig. 2). The citing paper counts in Ecosystem services, Landscapes & land use and Threatened species topics were 69-73% of predicted counts (Fig. 2). Protected areas and Human-wildlife interactions had few citing papers (55 and 51% of predicted counts, respectively), low even relative to the small body of core NEC papers in the topics (Fig. 2).

### Growth in NEC research over time

NEC research, including both core and citing papers, increased over time in all topics, rising from near-zero in 2006 (Fig. S2, Table S5). The rate of publication growth has slowed across all topics, particularly since 2020 (Fig. S2); over time, the number of new NEC papers published each year has levelled off (Fig. S2). Examining publication growth as a yearly rank among topics, in 2025 we found that eleven topics were within three ranks of their initial position in 2010 (Fig. 3). Five topics increased in ranking: Marine & coastal ecology increased from the 11th fastest growing topic to the fastest, Soil carbon & nutrients from 10th to 2nd, Conservation science from 7th to 3rd, Urban ecology from 12th to 6th and Regional studies from 20th to 15th (Fig. 3). Four topics decreased in ranking: Freshwater & wetlands decreased from 4th to 8th, Global change biology from 3rd to 10th, Ecological restoration from 5th to 13th and Landscapes & land use from 9th to 17th (Fig. 3).

**Figure 3:**
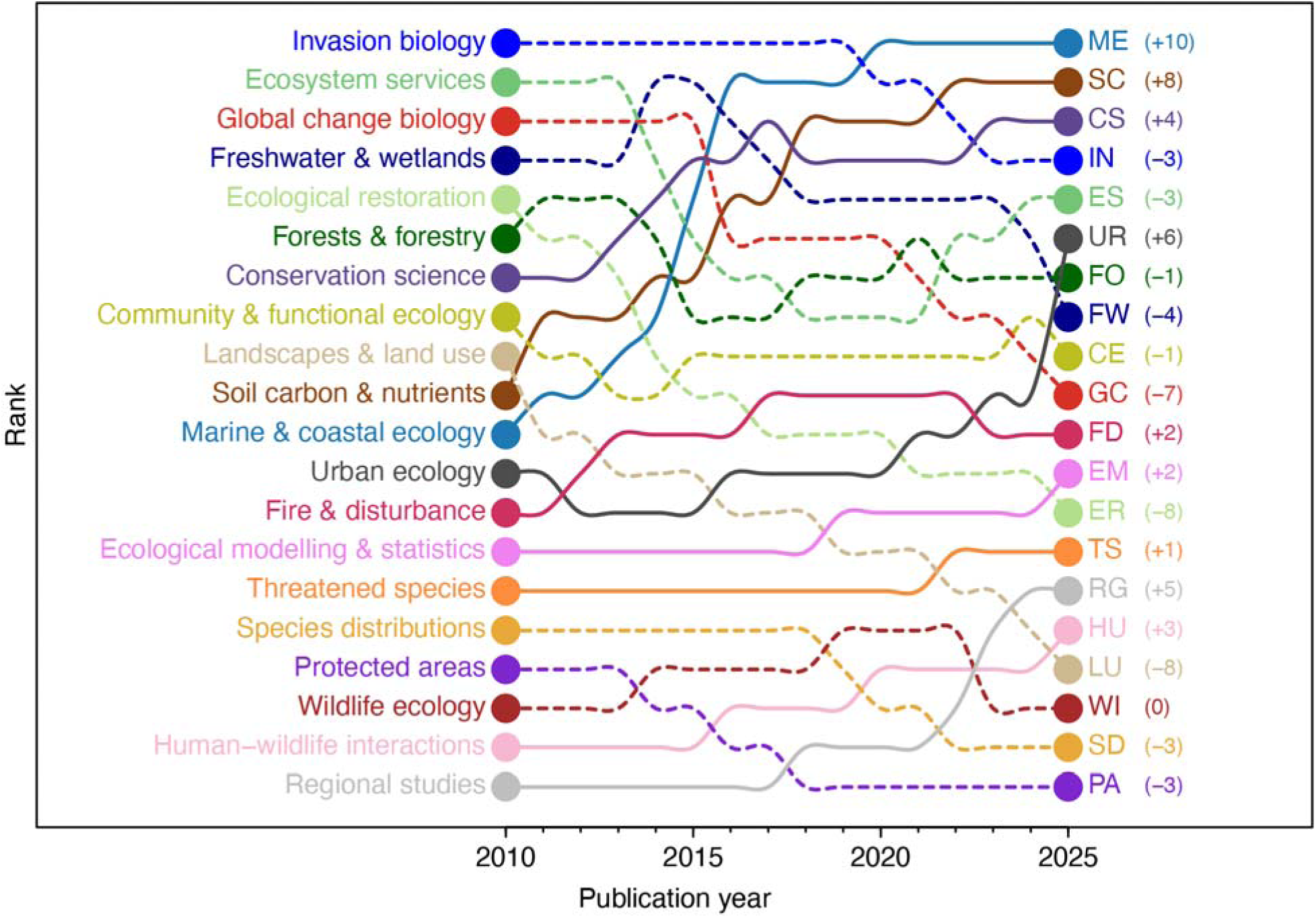
Rank of new Novel Ecosystem Concept (NEC) publications in applied ecology topics from 2010 to 2025 (includes core papers using the term “novel ecosystem” and papers citing core papers). For each year, higher ranked topics published more NEC papers than lower ranks. Ranks were established using a publication growth model (Fig. S2). Topics that increased in rank over time are shown with solid lines; topics with stable or decreasing ranks are shown with dashed lines. Numbers in parentheses indicate change in ranking between 2010 and 2025.

### Citation connectivity between NEC topics

Our citation model predicted the number of citations a “citing topic” made to a “cited topic”, using the core and citing NEC literature combined. We found citations increased with the number of core NEC papers in both the citing and cited topic (Fig. 4A). Larger topics in the applied ecology sample had a significant but near-zero effect on citation counts (Fig. 4B, Table S6). Topic size fixed effects captured just over half of the variation in citation counts (marginal R^2^ = 0.530). Linguistic similarity between citing and cited topics was a strong positive predictor of citation count (Fig. 4C, Table S7). Adding linguistic similarity to the model increased marginal R^2^ to 0.713, a 0.183 improvement over the size-correction model.

**Figure 4:**
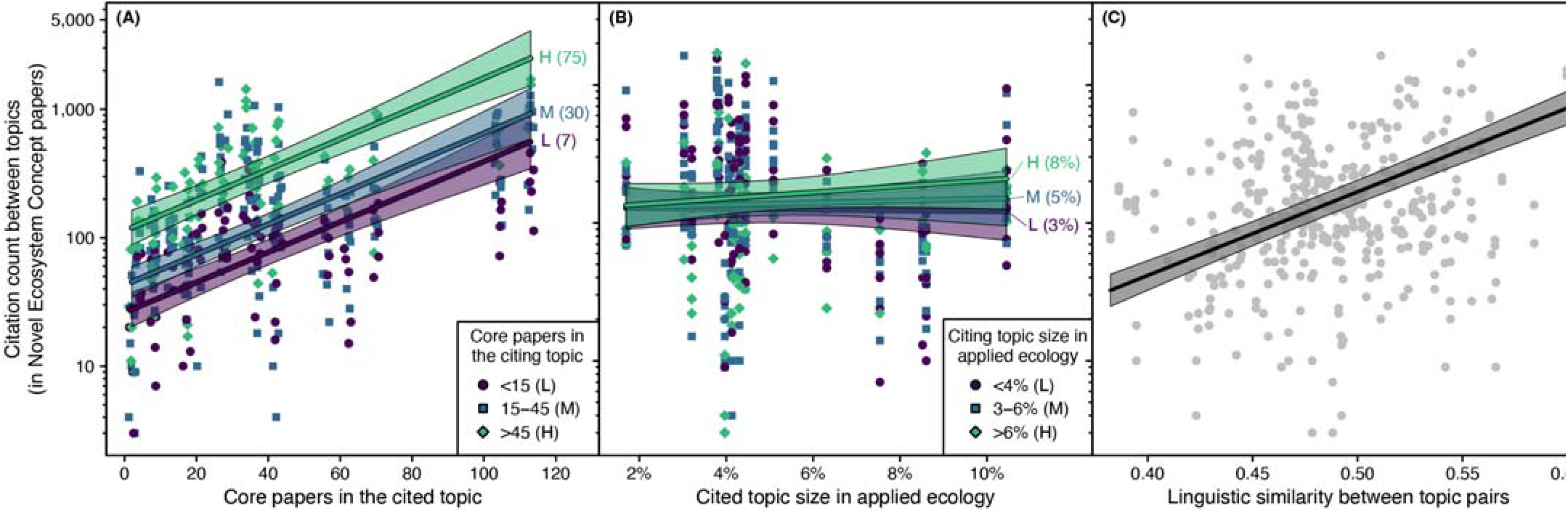
Citation count correlation between topic pairs and topic size and linguistic similarity fixed effects. Citations were counted between all core Novel Ecosystem Concept papers (NEC: using the term “novel ecosystem”) and papers citing them. **(A)** Citation counts were positively linked to the number of core NEC papers in both the cited (x-axis) and citing topic (point color and shape), **(B)** not strongly related to the overall size of citing and cited topic in a sample of applied ecology research and **(C)** positively correlated with linguistic similarity between topics. Linguistic similarity was estimated using the Bray-Curtis dissimilarity of paper-topic probabilities from the Latent Dirichlet Allocation topic model. Lines and shading reflect model predictions with associated 95% confidence intervals. For (A-B), model predictions were made at 10% (Low: L), mean (M) and 90% (High: H) quantiles of citing topic variables. These lines are partial regression slopes, holding all other fixed effects at their mean.

Citing and cited topic random intercepts explained an additional 15.5% of variation in citation counts (conditional R^2^ of 0.868 - marginal R^2^ of 0.713 in the linguistic similarity model). Positive citing and cited intercepts reflected “hub” topics that were central to the NEC literature. Global change biology, Ecological restoration and Species distributions fell into this quadrant (Fig. 5). Three topics had negative citing and cited random intercepts, which reflected “spokes” that were more weakly connected to the broader NEC literature: Human-wildlife interactions, Freshwater & wetlands and Soil carbon & nutrients (Fig. 5). Two topics were “scouts” with a positive citing intercept and a negative cited intercept: Marine & coastal ecology and Regional studies (Fig. 5). Finally, two topics, Ecosystem services and Protected areas, fell in the “authorities” quadrant. These topics had a negative citing intercept and a positive cited intercept, but both random effects were close to zero (Fig. 5).

**Figure 5:**
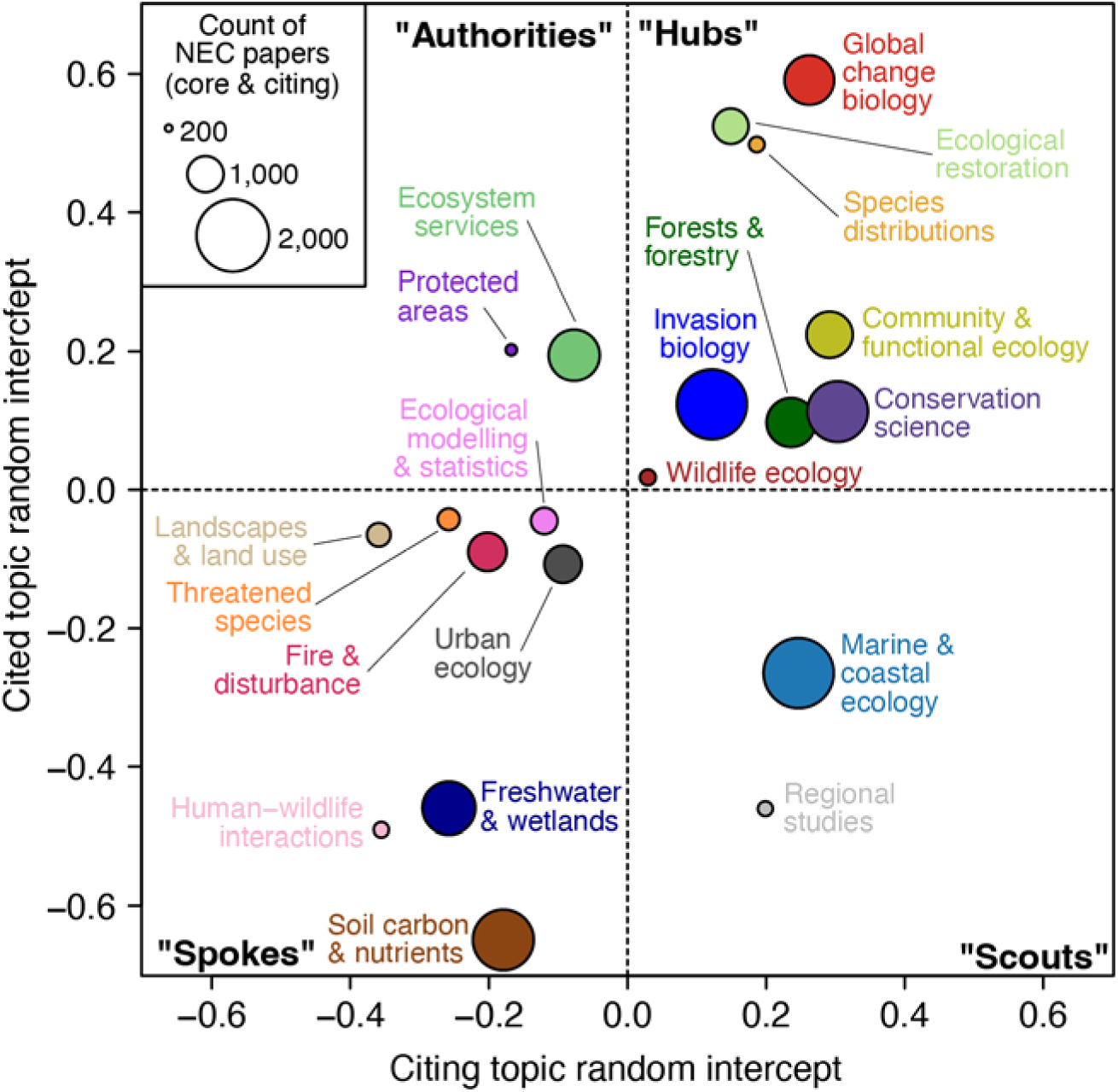
Random intercepts from citation count model for the Novel Ecosystem Concept (NEC) literature. X- and y-axes describe whether a topic tended to cite (“citing topic”) and be cited by (“cited topic”) other topics more (+) or less (-) than predicted from size and linguistic similarity fixed effects. Random intercept combination classes were “authorities” that were underciting and overcited, “hubs” that were overciting and overcited, “scouts” that were overciting and undercited and “spokes” that were underciting and undercited. Point sizes are scaled to the count of NEC core and citing papers (sum of x and y axes in Fig. 2).

## Discussion

The Novel Ecosystem Concept (NEC) promises a transformational lens to view ecosystem state and management under irreversible human-induced pressures. We examined the development of NEC research over the last twenty years, finding a complex and multidisciplinary literature. Research priorities have shifted away from some topics that dominated early NEC research, including Ecological restoration, Landscapes & land use and Global change biology and toward topic areas studying ecosystem-specific impacts: Freshwater & wetlands, Invasion biology, Marine & coastal ecology, Soil carbon & nutrients and Urban ecology. While the number of new NEC publications has grown each year, growth rates have slowed, and we identified some dominant areas of applied ecology (Protected areas, Human-wildlife interactions and Threatened species) where novel ecosystems have not been well-studied. In aggregate our results suggest the NEC is struggling to transition from a narrowly defined restoration framework to a broadly applied lens for ecosystem management in the Anthropocene. Despite this, our analysis highlights emerging NEC research areas in aquatic, soil and urban systems, with particularly rapid growth in the study of novel marine ecosystems. Our results highlight the power of bibliometric study to identify trends in ecological research and highlight opportunities to connect complimentary research agendas to improve on-the-ground ecosystem outcomes.

### Research topics and trajectories of the NEC

The NEC has been widely studied in applied ecology, with a publication record of more than 20 core papers in 13 of the 20 topics of applied ecology. This wide adoption reflects the value of the NEC to fill a conceptual gap in the study or management of human-altered systems, reflecting the near-universal human impact on ecosystems (Keck et al., 2025) that has been present for millennia (Ellis et al., 2021). The topic landscape of the NEC has shifted over time from restoration and land-management to topics dealing with severe and widespread human impacts in specific ecosystem types. The value of the NEC to these topics is clear. Outside of islands, removal of invasive species is often functionally impossible and intensive efforts produce only moderate success (Kettenring & Adams, 2011). Freshwater and marine systems are ecologically sensitive, often invaded by pernicious species and disproportionately impacted by landscape-scale resource use (Cooke et al., 2022) and climate change (Reid et al., 2019). Soil carbon & nutrients, alongside carbon storage research, included industrial reclamation and mine remediation, where even regaining basic ecosystem functioning is a core challenge (Rowland et al., 2009). The NEC, as a restoration triage and decision-making tool, has ready applicability to such degraded systems.

Applied ecology topics with low NEC engagement were strongly associated with conservation (Fig. S3). Early critiques of the NEC emerged from conservation biologists (Aronson et al., 2014; Kattan et al., 2016; Murcia et al., 2014). Our results indicate that this early resistance has persisted, despite the explicit adoption of conservation perspectives within NEC frameworks (Hobbs et al., 2009) and the growing recognition that conservation alone cannot arrest diversity loss from ecosystem degradation (Gann et al., 2026; Jones et al., 2018; Possingham et al., 2015). Deliberately managing novel ecosystems has moderate support as an option for biodiversity conservation (Clement et al., 2023), but this does not appear to be detectable in the NEC literature.

### Connectivity across the NEC

Despite the diffuse topic composition of the NEC, we could reliably predict citation counts between topic pairs using their linguistic similarity and size-based sampling effects, an indication that NEC research is well-connected overall. Most topics made and received citations in line with model expectations, with three exceptions: (1) Ecological restoration and Global change biology were core hubs of NEC citations, (2) Human-wildlife interactions, Freshwater & wetlands and Soil carbon & nutrients were relatively disconnected from the wider NEC literature, and (3) Marine & coastal ecology is emerging as a new core NEC research area.

The NEC literature was oriented around two major hub topics: Ecological restoration and Global change biology. These topics research the global and local processes that drive the formation of novel ecosystems (Hobbs et al., 2009). Both topics had low counts of core publications mentioning “novel ecosystems”. This suggests most of the engagement with hub topics is in the citing literature, using the perspective of the NEC to understand how climate change will impact management goals (Heller & Zavaleta, 2009) and influence restoration design (Harris et al., 2006; Simonson et al., 2021). While the NEC contributes to this discussion, our results suggest it is not a core component and, with declining publication growth ranking, both topics have become less central over recent decades.

Three topics were less connected to the broader NEC literature and may represent research barriers. Human-wildlife interactions had low rates of NEC publication as well as low citation connectivity. While the NEC has studied wildlife explicitly (e.g., Cadotte et al., 2021; Kennedy et al., 2018), it emerged from plant-dominated restoration research; this barrier may reflect the result of an entrenched plant-animal divide in biological research (Minelli, 2022). Freshwater & wetlands and Soil carbon & nutrients were not well-connected to the broader NEC literature, but this was not the case for other system-based topics (e.g., Urban ecology). Understanding land-water and soil-plant feedbacks are essential to successful ecosystem management (Rudgers et al., 2025; Wilcox, 2010), but our results indicate this importance has not translated into research and citation of the NEC.

Marine & coastal ecology rapidly grew to be the dominant NEC topic in 2025. Despite this growth and citation of marine NEC literature, the framework translating the NEC for marine ecosystems has only received minor attention (Schläppy & Hobbs, 2019). Instead, marine NEC research has studied the emergence of novel coral reefs states (Bulleri et al., 2020; Graham et al., 2014), ecological futures under climate change (Gillie et al., 2024) and non-traditional intervention methods using marine infrastructure (van Elden et al., 2022) and assisted evolution (van Oppen et al., 2015). Marine biologists appear to be engaging with the NEC as a lens to contextualize anthropogenic change rather than a specific management framework, aligning with efforts to broaden the NEC to describe “novel natures” (Montana et al., 2024). If this perspective can be integrated back into the wider literature, research in Marine & coastal ecology represents a valuable exploration and evolution of the NEC.

### Diagnosing the state of NEC research

While the number of applied ecology topics with some NEC research may be a sign of wide appeal, it may also indicate a “concept without a home”, lacking the centralization of other specialized areas of applied ecology. The NEC subtopic within Ecosystem services only comprised 38 papers, 5% of the 743 core NEC papers identified in our literature search. The diffuse and decelerating state of NEC research may be due to three factors. First, the NEC may simply be approaching the peak of its appeal, following the natural “steep rise, long tail” trajectory of research ideas (Tattershall et al., 2021). Alternatively, the NEC may not be relevant for much of applied ecology, due to either an inability to translate and operationalize the concept (Kattan et al., 2016), or a potential misalignment between the values in the NEC and those of applied ecologists (Thompson, 2017). Finally, it may be that the NEC has acted as a transitional concept, contributing to emerging frameworks of ecosystem management (e.g., Schuurman et al., 2022) rather than being adopted directly.

The NEC has struggled to define “novel” in a way that is easily applied to ecosystem management (Murcia et al., 2014; Truitt et al., 2015), even as the term “novel ecosystem” was initially intended to be descriptive (Larson, 2016). The NEC definition of novel ecosystems combines a classification of the current ecological state, historical human alteration and predicted resistance to future change, including potential management action (Hobbs et al., 2013). The complexity and challenge of satisfying this definition may have led to broad engagement with the concept of novel ecosystems rather than trying to directly operationalise underlying management framework. Our NEC literature dataset supports this conclusion; for every core paper that used the term “novel ecosystem”, we found 27 papers that cited the NEC literature without using the term. This peripheral engagement suggests that, currently, the NEC functions more as a theoretical framing device rather than an area of direct management value. Work to disentangle the facets of the novel ecosystem definition may provide a pathway forward. For example, “novelty”, a continuous measure of ecological difference, has been quantified in ecological (Finsinger et al., 2017) and abiotic contexts (Radeloff et al., 2015; Williams et al., 2007). Novelty as an ecological attribute is now formalized into frameworks for measurement (Staples et al., 2025) and identifying potential novel ecosystems (Kerr et al., 2026). There are further opportunities to relate ecological novelty to dynamic-systems theories of regime shift and alternative stable states (Mata et al., 2025), which formalize the thresholds and hysteresis that make some novel ecosystem changes difficult or impossible to reverse (e.g., Wolff et al., 2019). If the declining growth of the NEC is due to the challenge of defining “novel”, new quantitative perspectives and deeper integration with ecological theory may provide a pathway forward.

The challenge to NEC adoption may also be driven by conflicting values within ecological research. While scientists are taught to favor axiological neutrality, limiting how their intrinsic values impact their work, this is rarely true in practice (Weber, 1958). Ecologists are strongly informed by a passion for nature and desire to make a difference, values that shape our scientific lens (Cardou & Vellend, 2023). Many conservation biologists consider themselves advocates for species that cannot advocate for themselves (Brussard & Tull, 2007). The NEC, which espouses a practical approach to identifying and promoting the value in human-altered ecosystems, may be jarring to ecologists who value advocacy (Standish et al., 2013). The NEC explicitly “lowers the bar” for restoration by allowing for non-historical targets (Perring et al., 2013), which was critiqued for allowing accountability for ecological degradation to be avoided under the guise of calling the system “novel” (Murcia et al., 2014). Understanding and engaging with conflicting ecological values is key to ecosystem outcomes (Cardou & Vellend, 2023; Norberg et al., 2022), and our results suggest this conflict may be detectable within the ecological literature.

Finally, four approaches to ecosystem management reflect the NEC perspective in concept but not in name. The Resist-Accept-Direct framework orients management to changed conditions, allowing a choice of resisting changes via intervention, accepting the ecological consequences or directing the system along a more favorable trajectory (Millar et al., 2007; Williams, 2022).

Rewilding focuses on the restoration of ecological processes rather than the recovery of particular ecosystem states (Perino et al., 2019), an approach that overlaps with the explicit values-based nature of the NEC (Klop-Toker et al., 2020). Nature-based solutions aim to co-develop resilience and biodiversity, but these projects need to consider what is socially and financially acceptable as much as what is ecologically possible (Frantzeskaki et al., 2019). And even within Ecological restoration, a topic where NEC research has slowed, the Society for Ecological Restoration guidelines state that *“the reference model [as a restoration goal] is not necessarily the same as the historic state because all ecosystems (even those with high integrity) change over time”* (Gann et al., 2026). This is, in effect, recognition that novel ecosystems may be legitimate restoration targets, without explicitly referring to the NEC. While our results indicate research into “novel ecosystems” is slowing, broad research and citation across the topics of applied ecology suggest the basic tenets of the NEC have been widely adapted and adopted.

### Concluding remarks

Recognition of how past human actions have co-produced the ecosystems of the present is growing (Ellis et al., 2021). In a rapidly changing world, our science needs to be adaptive and reflective, drawing from all relevant knowledge to improve ecological outcomes (Cadotte et al., 2017). Despite growth slowing over time, the NEC has been studied and cited by almost all topics of applied ecology, including rapidly advancing marine research that seeks to promote ecological resilience in the Anthropocene (Bulleri et al., 2020). The oceans may be a catalyst to creating an NEC Renaissance. The concept of novelty is being quantified into measurements of ecological change and has been adopted in a variety of contexts (Kerr et al., 2026; Staples et al., 2025), and the perspective of the NEC has been integrated into new management and restoration frameworks. The Anthropocene promises climate regimes and human pressures that will irreversibly alter the biosphere (Rapacciuolo et al., 2019). The NEC has laid the groundwork for describing and managing this change and there are promising signs that this impact will continue to deliver valuable social-ecological outcomes into the future.

## Author contributions

Conception, design, data acquisition, analysis and writing was led by TLS, with substantial intellectual contribution across all areas by coauthors. This study was an analysis of English academic literature, with a globally distributed authorship team. Our study fails to include non-English research and authors from several major regions including Africa and Asia, a caveat we aim to address in future research.

## Acknowledgements

We are grateful to Clarivate™ for collating and providing access to publication metadata via their Web of Science™ platform (© Clarivate 2026). All rights reserved). This work was facilitated by the NovelEco Symposium, which is funded by the European Research Council (ERC) under the European Union’s Horizon 2020 Research & Innovation programme (Grant Agreement No. 101002440). TLS was supported by an Australian Research Council Discovery Early Career Research Award (DE240100398). AO, MRK and JCS were supported by Center for Ecological Dynamics in a Novel Biosphere (ECONOVO), funded by Danish National Research Foundation (grant DNRF173). SLM was supported by the FWO research grant 1203025N. YMB was supported through financial support of Research Ireland, Northern Ireland’s Department of Agriculture, Environment and Rural Affairs (DAERA), UK Research and Innovation (UKRI) via the International Science Partnerships Fund (ISPF) under Grant number [22/CC/11103] at the Co-Centre for Climate + Biodiversity + Water.

## Data availability

Data and R scripts necessary to reproduce all analyses are available at https://github.com/TimothyStaples/novelEcoCitationMapping. This code will be versioned in Zenodo and cited with a DOI number upon manuscript acceptance.

## Reference list

Anthropic. (2026). *Claude Sonnet 4.6* (Sonnet 4.6). Anthropic. https://claude.ai

Araújo, M. B., Cabeza, M., Thuiller, W., Hannah, L., & Williams, P. H. (2004). Would climate change drive species out of reserves? An assessment of existing reserve-selection methods. Global Change Biology, 10(9), 1618–1626. 10.1111/j.1365-2486.2004.00828.x

Aronson, J., Murcia, C., Kattan, G. H., Moreno-Mateos, D., Dixon, K., & Simberloff, D. (2014). The road to confusion is paved with novel ecosystem labels: A reply to Hobbs et al. Trends in Ecology and Evolution, 29(12), 646–647. 10.1016/j.tree.2014.09.011

Bornmann, L., & Daniel, H. D. (2008). What do citation counts measure? A review of studies on citing behavior. Journal of Documentation, 64(1), 45–80. 10.1108/00220410810844150

Brooks, M. E., Kristensen, K., van Benthem, K. J., Magnusson, A., Berg, C. W., Nielsen, A., Skaug, H. J., Maechler, M., & Bolker, B. M. (2017). glmmTMB Balances Speed and Flexibility Among Packages for Zero-inflated Generalized Linear Mixed Modeling. The R Journal, 9(2), 378–400. 10.32614/RJ-2017-066

Brudvig, L. A., Barak, R. S., Bauer, J. T., Caughlin, T. T., Laughlin, D. C., Larios, L., Matthews, J. W., Stuble, K. L., Turley, N. E., & Zirbel, C. R. (2017). Interpreting variation to advance predictive restoration science. Journal of Applied Ecology, 54(4), 1018–1027. 10.1111/1365-2664.12938

Brussard, P. F., & Tull, J. C. (2007). Conservation Biology and Four Types of Advocacy. Conservation Biology, 21(1), 21–24. http://www.jstor.org/stable/4124637

Bulleri, F., Batten, S., Connell, S. D., Benedetti-Cecchi, L., Gibbons, M., Nugues, M., & Gribben, P. (2020). Human pressures and the emergence of novel marine ecosystems. In S. J. Hawkins, A. L. Allcock, A. E. Bates, L. B. Firth, I. P. Smith, S. E. Swearer, A. J. Evans, P. A. Todd, B. D. Russell, & C. D. McQuaid (Eds.), Oceanography and Marine Biology: An Annual Review, Volume 58 (pp. 441–493). Taylor & Francis. 10.1201/9780429351495-9

Cadotte, M. W., Barlow, J., Nuñez, M. A., Pettorelli, N., & Stephens, P. A. (2017). Solving environmental problems in the Anthropocene: the need to bring novel theoretical advances into the applied ecology fold. Journal of Applied Ecology, 54(1), 1–6. 10.1111/1365-2664.12855

Cadotte, M. W., Potgieter, L. J., Wang, C. J., & MacIvor, J. S. (2021). Invasion theory as a management tool for increasing native biodiversity in urban ecosystems. Journal of Applied Ecology, 58(11), 2394–2403. 10.1111/1365-2664.13953

Cardou, F., & Vellend, M. (2023). Stealth advocacy in ecology and conservation biology. Biological Conservation, 280, 109968. 10.1016/j.biocon.2023.109968

Clement, S., Standish, R. J., & Kennedy, P. L. (2023). Expert preferences on options for biodiversity conservation under climate change. Global Environmental Change, 83, 102759. 10.1016/j.gloenvcha.2023.102759

Dias, L., Gerlach, M., Scharloth, J., & Altmann, E. G. (2018). Using text analysis to quantify the similarity and evolution of scientific disciplines. Royal Society Open Science, 5(1), 171545. 10.1098/rsos.171545

Dombrovski, V. C., Zhurauliou, D. V, & Ashton-Butt, A. (2022). Long-term effects of rewilding on species composition: 22 years of raptor monitoring in the Chernobyl Exclusion Zone. Restoration Ecology, 30(8), e13633. 10.1111/rec.13633

Ellis, E. C. (2021). Land Use and Ecological Change: A 12,000-Year History. Annual Review of Environment and Resources, 46(Volume 46, 2021), 1–33. 10.1146/annurev-environ-012220-010822

Ellis, E. C., Gauthier, N., Klein Goldewijk, K., Bliege Bird, R., Boivin, N., Díaz, S., Fuller, D. Q., Gill, J. L., Kaplan, J. O., Kingston, N., Locke, H., McMichael, C. N. H., Ranco, D., Rick, T. C., Shaw, M. R., Stephens, L., Svenning, J.-C., & Watson, J. E. M. (2021). People have shaped most of terrestrial nature for at least 12,000 years. Proceedings of the National Academy of Sciences, 118(17), e2023483118–e2023483118. 10.1073/pnas.2023483118

Ellis, E. C., Klein Goldewijk, K., Siebert, S., Lightman, D., & Ramankutty, N. (2010). Anthropogenic transformation of the biomes, 1700 to 2000. Global Ecology and Biogeography, 19(5), 589–606. 10.1111/j.1466-8238.2010.00540.x

Evers, C. R., Wardropper, C. B., Branoff, B., Granek, E. F., Hirsch, S. L., Link, T. E., Olivero-Lora, S., & Wilson, C. (2018). The ecosystem services and biodiversity of novel ecosystems: A literature review. Global Ecology and Conservation, 13, e00362–e00362. 10.1016/j.gecco.2017.e00362

Finsinger, W., Giesecke, T., Brewer, S., & Leydet, M. (2017). Emergence patterns of novelty in European vegetation assemblages over the past 15 000 years. Ecology Letters, 20(3), 336–346. 10.1111/ele.12731

Frantzeskaki, N., McPhearson, T., Collier, M. J., Kendal, D., Bulkeley, H., Dumitru, A., Walsh, C., Noble, K., van Wyk, E., Ordóñez, C., Oke, C., & Pintér, L. (2019). Nature-Based Solutions for Urban Climate Change Adaptation: Linking Science, Policy, and Practice Communities for Evidence-Based Decision-Making. BioScience, 69(6), 455–466. 10.1093/biosci/biz042

Gann, G. D., McDonald, T., Walder, B., Aronson, J., Nelson, C. R., Jonson, J., Hallett, J. G., Eisenberg, C., Guariguata, M. R., Liu, J., Hua, F., Echeverría, C., Gonzales, E., Shaw, N., Decleer, K., & Dixon, K. W. (2019). International principles and standards for the practice of ecological restoration. Second edition. Restoration Ecology, 27(S1), S1–S46. 10.1111/rec.13035

Gillie, E. R., Bryndum-Buchholz, A., Willis, S. G., & Eddy, T. D. (2024). Exploring novel North Water Polynya ecosystems under climate change. PLOS Climate, 3(10), e0000490-. 10.1371/journal.pclm.0000490

Graham, N. A. J., Cinner, J. E., Norström, A. V, & Nyström, M. (2014). Coral reefs as novel ecosystems: embracing new futures. Current Opinion in Environmental Sustainability, 7, 9–14. 10.1016/j.cosust.2013.11.023

Hallett, L. M., Diver, S., Eitzel, M. V, Olson, J. J., Ramage, B. S., Sardinas, H., Statman-Weil, Z., & Suding, K. N. (2013). Do we practice what we preach? Goal setting for ecological restoration. Restoration Ecology, 21(3), 312–319. 10.1111/rec.12007

Harris, J. A., Hobbs, R. J., Higgs, E., & Aronson, J. (2006). Ecological restoration and global climate change. Restoration Ecology, 14(2), 170–176. 10.1111/j.1526-100X.2006.00136.x

Heller, N. E., & Zavaleta, E. S. (2009). Biodiversity management in the face of climate change: A review of 22 years of recommendations. Biological Conservation, 142(1), 14–32. 10.1016/j.biocon.2008.10.006

Hobbs, R. J. (2017). Novel ecosystems: Can’t we just pretend they’re not there? In Effective Conservation Science: Data Not Dogma (pp. 45–50). Oxford University Press. 10.1093/oso/9780198808978.003.0007

Hobbs, R. J., Arico, S., Aronson, J., Baron, J. S., Bridgewater, P., Cramer, V. A., Epstein, P. R., Ewel, J. J., Klink, C. A., Lugo, A. E., Norton, D., Ojima, D., Richardson, D. M., Sanderson, E. W., Valladares, F., Vilà, M., Zamora, R., & Zobel, M. (2006a). Novel ecosystems: Theoretical and management aspects of the new ecological world order. Global Ecology and Biogeography, 15(1), 1–7. 10.1111/j.1466-822X.2006.00212.x

Hobbs, R. J., Arico, S., Aronson, J., Baron, J. S., Bridgewater, P., Cramer, V. A., Epstein, P. R., Ewel, J. J., Klink, C. A., Lugo, A. E., Norton, D., Ojima, D., Richardson, D. M., Sanderson, E. W., Valladares, F., Vilà, M., Zamora, R., & Zobel, M. (2006b). Novel ecosystems: Theoretical and management aspects of the new ecological world order. Global Ecology and Biogeography, 15(1), 1–7. 10.1111/j.1466-822X.2006.00212.x

Hobbs, R. J., Hallett, L. M., Ehrlich, P. R., & Mooney, H. A. (2011). Intervention Ecology: Applying Ecological Science in the Twenty-first Century. BioScience, 61(6), 442–450. 10.1525/bio.2011.61.6.6

Hobbs, R. J., Higgs, E., Hall, C. M., Bridgewater, P., Chapin, F. S., Ellis, E. C., Ewel, J. J., Hallett, L. M., Harris, J., Hulvey, K. B., Jackson, S. T., Kennedy, P. L., Kueffer, C., Lach, L., Lantz, T. C., Lugo, A. E., Mascaro, J., Murphy, S. D., Nelson, C. R., … Yung, L. (2014a). Managing the whole landscape: Historical, hybrid, and novel ecosystems. Frontiers in Ecology and the Environment, 12(10), 557–564. 10.1890/130300

Hobbs, R. J., Higgs, E., Hall, C. M., Bridgewater, P., Chapin, F. S., Ellis, E. C., Ewel, J. J., Hallett, L. M., Harris, J., Hulvey, K. B., Jackson, S. T., Kennedy, P. L., Kueffer, C., Lach, L., Lantz, T. C., Lugo, A. E., Mascaro, J., Murphy, S. D., Nelson, C. R., … Yung, L. (2014b). Managing the whole landscape: Historical, hybrid, and novel ecosystems. Frontiers in Ecology and the Environment, 12(10), 557–564. 10.1890/130300

Hobbs, R. J., Higgs, E., & Harris, J. A. (2009a). Novel ecosystems: implications for conservation and restoration. Trends in Ecology and Evolution, 24(11), 599–605. 10.1016/j.tree.2009.05.012

Hobbs, R. J., Higgs, E., & Harris, J. A. (2009b). Novel ecosystems: implications for conservation and restoration. Trends in Ecology and Evolution, 24(11), 599–605. 10.1016/j.tree.2009.05.012

Hobbs, R. J., Higgs, E. S., & Hall, C. M. (2013). Defining Novel Ecosystems. In Novel Ecosystems: Intervening in the New Ecological World Order (pp. 58–60). 10.1002/9781118354186.ch6

Hulvey, K. B., Standish, R. J., Hallett, L. M., Starzomski, B. M., Murphy, S. D., Nelson, C. R., Gardener, M. R., Kennedy, P. L., Seastedt, T. R., & Suding, K. N. (2013). Incorporating Novel Ecosystems into Management Frameworks. In Novel Ecosystems (pp. 157–171). 10.1002/9781118354186.ch18

Jackson, S. T., & Hobbs, R. J. (2009). Ecological Restoration in the Light of Ecological History. Science, 325(5940), 567–569. 10.1126/science.1172977

Jagodziński, A. M., Wierzcholska, S., Dyderski, M. K., Horodecki, P., Rusińska, A., Gdula, A. K., & Kasprowicz, M. (2018). Tree species effects on bryophyte guilds on a reclaimed post-mining site. Ecological Engineering, 110, 117–127. 10.1016/j.ecoleng.2017.10.015

Jaureguiberry, P., Titeux, N., Wiemers, M., Bowler, D. E., Coscieme, L., Golden, A. S., Guerra, C. A., Jacob, U., Takahashi, Y., Settele, J., Díaz, S., Molnár, Z., & Purvis, A. (2026). The direct drivers of recent global anthropogenic biodiversity loss. Science Advances, 8(45), eabm9982. 10.1126/sciadv.abm9982

Jelodar, H., Wang, Y., Yuan, C., Feng, X., Jiang, X., Li, Y., & Zhao, L. (2019). Latent Dirichlet allocation (LDA) and topic modeling: models, applications, a survey. Multimedia Tools and Applications, 78(11), 15169–15211. 10.1007/s11042-018-6894-4

Jones, H. P., Jones, P. C., Barbier, E. B., Blackburn, R. C., Rey Benayas, J. M., Holl, K. D., McCrackin, M., Meli, P., Montoya, D., & Mateos, D. M. (2018a). Restoration and repair of Earth’s damaged ecosystems. Proceedings of the Royal Society B: Biological Sciences, 285(1873). 10.1098/rspb.2017.2577

Jones, H. P., Jones, P. C., Barbier, E. B., Blackburn, R. C., Rey Benayas, J. M., Holl, K. D., McCrackin, M., Meli, P., Montoya, D., & Mateos, D. M. (2018b). Restoration and repair of Earth’s damaged ecosystems. Proceedings of the Royal Society B: Biological Sciences, 285(1873), 20172577. 10.1098/rspb.2017.2577

Kattan, G. H., Aronson, J., & Murcia, C. (2016). Does the novel ecosystem concept provide a framework for practical applications and a path forward? A reply to Miller and Bestelmeyer. Restoration Ecology, 24(6), 714–716. 10.1111/rec.12453

Keck, F., Peller, T., Alther, R., Barouillet, C., Blackman, R., Capo, E., Chonova, T., Couton, M., Fehlinger, L., Kirschner, D., Knüsel, M., Muneret, L., Oester, R., Tapolczai, K., Zhang, H., & Altermatt, F. (2025). The global human impact on biodiversity. Nature, 641(8062), 395–400. 10.1038/s41586-025-08752-2

Kennedy, P. L., Fontaine, J. B., Hobbs, R. J., Johnson, T. N., Boyle, R., & Lueders, A. S. (2018). Do novel ecosystems provide habitat value for wildlife? Revisiting the physiognomy vs. floristics debate. Ecosphere, 9(3), e02172. 10.1002/ecs2.2172

Kerr, M. R., Svenning, J.-C., Riede, F., Buitenwerf, R., Abraham, A. J., Marjakangas, E.-L., Sykut, M., & Ordonez, A. (2026). The making of novel ecosystems: A process-based framework for measurement, analysis and application. Methods in Ecology and Evolution, 17(3), 683–704. 10.1111/2041-210x.70255

Klop-Toker, K., Clulow, S., Shuttleworth, C., & Hayward, M. W. (2020). Are novel ecosystems the only novelty of rewilding? Restoration Ecology, 28(6), 1318–1320. 10.1111/rec.13241

Kreyling, J., Tanneberger, F., Jansen, F., van der Linden, S., Aggenbach, C., Blüml, V., Couwenberg, J., Emsens, W.-J., Joosten, H., Klimkowska, A., Kotowski, W., Kozub, L., Lennartz, B., Liczner, Y., Liu, H., Michaelis, D., Oehmke, C., Parakenings, K., Pleyl, E., … Jurasinski, G. (2021). Rewetting does not return drained fen peatlands to their old selves. Nature Communications, 12(1), 5693. 10.1038/s41467-021-25619-y

Larson, B. M. H. (2016). New Wine and Old Wineskins? Novel Ecosystems and Conceptual Change. Nature and Culture, 11(2), 148–164. 10.3167/nc.2016.110202

Lawler, J. J. (2009). Climate Change Adaptation Strategies for Resource Management and Conservation Planning. In R. S. Ostfeld & W. H. Schlesinger (Eds.), YEAR IN ECOLOGY AND CONSERVATION BIOLOGY 2009 (Vol. 1162). 10.1111/j.1749-6632.2009.04147.x

Lindenmayer, D. B., Fischer, J., Felton, A., Crane, M., Michael, D., Macgregor, C., Montague-Drake, R., Manning, A., & Hobbs, R. J. (2008). Novel ecosystems resulting from landscape transformation create dilemmas for modern conservation practice. Conservation Letters, 1(3), 129–135. 10.1111/j.1755-263x.2008.00021.x

Maes, S. L., Perring, M. P., Cohen, R., Akinnifesi, F. K., Bargués-Tobella, A., Bastin, J.-F., Bauters, M., Bernardino, P. N., Brancalion, P. H. S., Bullock, J. M., Ellison, D., Fayolle, A., Fremout, T., Gann, G. D., Hishe, H., Holmgren, M., Ilstedt, U., Mahy, G., Messier, C., … Muys, B. (2024). Explore before you restore: Incorporating complex systems thinking in ecosystem restoration. Journal of Applied Ecology, 61(5), 922–939. 10.1111/1365-2664.14614

Mascaro, J., Hughes, R. F., & Schnitzer, S. A. (2012). Novel forests maintain ecosystem processes after the decline of native tree species. Ecological Monographs, 82(2), 221–228. 10.1890/11-1014.1

McNellie, M. J., Oliver, I., Dorrough, J., Ferrier, S., Newell, G., & Gibbons, P. (2020). Reference state and benchmark concepts for better biodiversity conservation in contemporary ecosystems. Global Change Biology, 26(12), 6702–6714. 10.1111/gcb.15383

Millar, C. I., Stephenson, N. L., & Stephens, S. L. (2007). Climate change and forests of the future: Managing in the face of uncertainty. Ecological Applications, 17(8), 2145–2151. 10.1890/06-1715.1

Minelli, A. (2022). Two-way exchanges between animal and plant biology, with focus on evo-devo. Frontiers in Ecology and Evolution, Volume 10-2022. 10.3389/fevo.2022.1057355

Montana, J., Heger, T., Kelz, R., Bischoff, A., Buitenwerf, R., Eser, U., Kung, K. T., Sattler, J., Schweiger, A. H., Searle, A., Teixeira, L. H., Travassos-Britto, B., & Higgs, E. (2024). From novel ecosystems to novel natures. Gaia - Ecological Perspectives for Science and Society, 33(1), 146–151. 10.14512/gaia.33.1.6

Murcia, C., Aronson, J., Kattan, G. H., Moreno-Mateos, D., Dixon, K., & Simberloff, D. (2014). A critique of the “novel ecosystem” concept. Trends in Ecology and Evolution, 29(10), 548–553. 10.1016/j.tree.2014.07.006

Norberg, J., Blenckner, T., Cornell, S. E., Petchey, O. L., & Hillebrand, H. (2022). Failures to disagree are essential for environmental science to effectively influence policy development. Ecology Letters, 25(5), 1075–1093. 10.1111/ele.13984

Perino, A., Pereira, H. M., Navarro, L. M., Fernández, N., Bullock, J. M., Ceau u, S., Cortés-Avizanda, A., van Klink, R., Kuemmerle, T., Lomba, A., Pe’er, G., Plieninger, T., Rey Benayas, J. M., Sandom, C. J., Svenning, J.-C., & Wheeler, H. C. (2019). Rewilding complex ecosystems. Science, 364(6438), eaav5570. 10.1126/science.aav5570

Perring, M. P., Standish, R. J., & Hobbs, R. J. (2013). Incorporating novelty and novel ecosystems into restoration planning and practice in the 21st century. Ecological Processes, 2(1), 18. 10.1186/2192-1709-2-18

Possingham, H. P., Bode, M., & Klein, C. J. (2015). Optimal Conservation Outcomes Require Both Restoration and Protection. PLOS Biology, 13(1), e1002052-. 10.1371/journal.pbio.1002052

Radeloff, V. C., Williams, J. W., Bateman, B. L., Burke, K. D., Carter, S. K., Childress, E. S., Cromwell, K. J., Gratton, C., Hasley, A. O., Kraemer, B. M., Latzka, A. W., Marin-Spiotta, E., Meine, C. D., Munoz, S. E., Neeson, T. M., Pidgeon, A. M., Rissman, A. R., Rivera, R. J., Szymanski, L. M., & Usinowicz, J. (2015). The rise of novelty in ecosystems. Ecological Applications, 25(8), 2051–2068. 10.1890/14-1781.1

Rapacciuolo, G., Rominger, A. J., Morueta-Holme, N., & Blois, J. L. (2019). Editorial: Ecological Non-equilibrium in the Anthropocene. Frontiers in Ecology and Evolution, Volume 7-2019. https://www.frontiersin.org/journals/ecology-and-evolution/articles/10.3389/fevo.2019.00428

Reid, A. J., Carlson, A. K., Creed, I. F., Eliason, E. J., Gell, P. A., Johnson, P. T. J., Kidd, K. A., MacCormack, T. J., Olden, J. D., Ormerod, S. J., Smol, J. P., Taylor, W. W., Tockner, K., Vermaire, J. C., Dudgeon, D., & Cooke, S. J. (2019). Emerging threats and persistent conservation challenges for freshwater biodiversity. Biological Reviews, 94(3), 849–873. 10.1111/brv.12480

Rowland, S. M., Prescott, C. E., Grayston, S. J., Quideau, S. A., & Bradfield, G. E. (2009). Recreating a Functioning Forest Soil in Reclaimed Oil Sands in Northern Alberta: An Approach for Measuring Success in Ecological Restoration. Journal of Environmental Quality, 38(4), 1580–1590. 10.2134/jeq2008.0317

Rudgers, J. A., Gehring, C. A., Taylor, D. L., Taylor, M. D., & Chung, Y. A. (2025). Integration of plant–soil feedbacks with resilience theory for climate change. Trends in Ecology & Evolution, 40(8), 749–759. 10.1016/j.tree.2025.05.001

Sandbrook, C. (2015). What is conservation? Oryx, 49(4), 565–566. 10.1017/S0030605315000952

Schuurman, G. W., Cole, D. N., Cravens, A. E., Covington, S., Crausbay, S. D., Hoffman, C. H., Lawrence, D. J., Magness, D. R., Morton, J. M., Nelson, E. A., & O’Malley, R. (2022). Navigating Ecological Transformation: Resist–Accept–Direct as a Path to a New Resource Management Paradigm. BioScience, 72(1), 16–29. 10.1093/biosci/biab067

Simonson, W. D., Miller, E., Jones, A., García-Rangel, S., Thornton, H., & McOwen, C. (2021). Enhancing climate change resilience of ecological restoration — A framework for action. Perspectives in Ecology and Conservation, 19(3), 300–310. 10.1016/j.pecon.2021.05.002

Staples, T. L., Blois, J., Cramer, K. L., Cunningham, E. T., Dornelas, M., Haberle, S. G., Heger, T., Kiessling, W., Magurran, A. E., O’Dea, A., Penny, A. M., Radeloff, V. C., Smith, J. A., Thuiller, W., Williams, J. W., & Pandolfi, J. M. (2025). A Conceptual Framework for Measuring Ecological Novelty. Global Ecology and Biogeography, 34(2), e70005–e70005. 10.1111/geb.70005

Staples, T. L., Dwyer, J. M., Wainwright, C. E., & Mayfield, M. M. (2019). Applied ecological research is on the rise but connectivity barriers persist between four major subfields. Journal of Applied Ecology, 56(6), 1492–1498. 10.1111/1365-2664.13373

Tattershall, E., Nenadic, G., & Stevens, R. D. (2021). Modelling trend life cycles in scientific research using the Logistic and Gompertz equations. Scientometrics, 126(11), 9113–9132. 10.1007/s11192-021-04137-0

Thompson, A. R. H. (2017). Managing for Virtue: The Moral Debate over Novel Ecosystems. *Worldviews: Global Religions*, Culture, and Ecology, 21(2), 134–151. 10.1163/15685357-02102003

Truitt, A. M., Granek, E. F., Duveneck, M. J., Goldsmith, K. A., Jordan, M. P., & Yazzie, K. C. (2015). What is Novel About Novel Ecosystems: Managing Change in an Ever-Changing World. Environmental Management, 55(6), 1217–1226. 10.1007/s00267-015-0465-5

United Nations General Assembly. (2010). Resolution 65/161: United Nations Decade on Biodiversity, 2011–2020. https://www.cbd.int/undb/goals/undb-unresolution.pdf

United Nations General Assembly. (2019). Resolution 73/284: United Nations Decade on Ecosystem Restoration (2021–2030). https://digitallibrary.un.org/record/3794317

van Elden, S., Meeuwig, J. J., & Hobbs, R. J. (2022). Offshore platforms as novel ecosystems: A case study from Australia’s Northwest Shelf. Ecology and Evolution, 12(2), e8496. 10.1002/ece3.8496

van Oppen, M. J. H., Oliver, J. K., Putnam, H. M., & Gates, R. D. (2015). Building coral reef resilience through assisted evolution. Proceedings of the National Academy of Sciences, 112(8), 2307–2313. 10.1073/pnas.1422301112

Weber, M. (1958). Science as a Vocation. Daedalus, 87(1), 111–134. http://www.jstor.org/stable/20026431

Wilcox, B. P. (2010). Transformative ecosystem change and ecohydrology: ushering in a new era for watershed management. Ecohydrology, 3(1), 126–130. 10.1002/eco.104

Williams, B. K., & Brown, E. D. (2024). Managing ecosystems with resist–accept–direct (RAD). Methods in Ecology and Evolution, 15(5), 796–805. 10.1111/2041-210X.14309

Williams, J., Jackson, S. T., & Kutzbach, J. E. (2007). Projected distributions of novel and disappearing climates by 2100 AD. Proceedings of the National Academy of Sciences of the United States of America, 104(14), 5738–5742. 10.1073/pnas.0606292104

Williams, J. W. (2022). RAD: A Paradigm, Shifting. BioScience, 72(1), 13–15. 10.1093/biosci/biab123

Williams, J. W., & Jackson, S. T. (2007). Novel climates, no-analog communities, and ecological surprises. Frontiers in Ecology and the Environment, 5(9), 475–482. 10.1890/070037

Williams, J. W., Ordonez, A., & Svenning, J.-C. (2021). A unifying framework for studying and managing climate-driven rates of ecological change. Nature Ecology & Evolution, 5(1), 17–26. 10.1038/s41559-020-01344-5

Zhang, Y., Chen, H., Lu, J., & Zhang, G. (2017). Detecting and predicting the topic change of Knowledge-based Systems: A topic-based bibliometric analysis from 1991 to 2016. Knowledge-Based Systems, 133, 255–268. 10.1016/j.knosys.2017.07.0

