## Supplementary Material for "Two decades of Novel Ecosystems: Literature analysis of topics, trends and connectivity"

Supplementary methods

*Querying Large Language Model for topic association names*

Claude Sonnet 4.6 was queried with the following prompt:

***"****I am going to give you a .csv file of stemmed keywords from a topic association model of ecological research. Rows reflect terms ranked by their importance to the topic and columns reflect each topic. I want you to provide a table with a short topic name (<30 characters) and an associated colour, aiming for contrast in data visualisation."*

This prompt was followed by the topic-associated terms in Table S1, provided in row-column format with column headings of "Topic1" to "Topic20". Claude output was scrutinized and modified by authors using their understanding of the ecological literature.

Supplementary Figures and Tables

***Table S1:*** *Topics of applied ecology as identified using a Latent Dirichlet Allocation model run on the titles, keywords and abstracts of each paper. Core papers used the term "novel ecosystem", citing papers cited at least one core paper and applied ecology sample reflect count from a general paper search to capture topics that did not contain substantial Novel Ecosystem Concept literature. Topic terms were the top 20 terms used more within a topic than across topics. Topic terms and example papers were used to name each topic.*

| **Topic name** | **Abbrev** | **Core paper count** | **Citing paper count** | **Applied ecology**  **sample count** | **Topic-associated terms** | **Example paper** |
| --- | --- | --- | --- | --- | --- | --- |
| Community & functional ecology | CE | 36 | 1240 | 1078 | communiti, divers, speci, function, rich, trait, composit, abund, differ, structur, group, assemblag, biodivers, site, gradient, across, environment, pattern, relat, taxonom | Baumberger, Teddy (2012) Plant community changes as ecological indicator of seabird colonies' impacts on Mediterranean Islands |
| Conservation science | CS | 70 | 1605 | 2215 | conserv, research, social, natur, knowledg, scienc, polici, valu, local, environment, public, govern, peopl, cultur, practic, stakehold, human, communiti, percept, scientif | Burgess, H. K. (2017) The science of citizen science: Exploring barriers to use as a primary research tool |
| Ecological modelling & statistics | EM | 13 | 723 | 884 | use, data, assess, method, indic, monitor, base, estim, analysi, map, studi, detect, inform, approach, sampl, measur, remot, survey, sens, evalu | Rasanen, Aleksi (2019) Data and resolution requirements in mapping vegetation in spatially heterogeneous landscapes |
| Ecological restoration | RE | 34 | 945 | 943 | restor, manag, strategi, approach, effect, decis, project, cost, plan, implement, develop, improv, target, benefit, success, requir, evalu, risk, action, design | Rodrigues, Ricardo R. (2009) On the restoration of high diversity forests: 30 years of experience in the Brazilian Atlantic Forest |
| Ecosystem services | ES | 113 | 1304 | 803 | ecosystem, ecolog, servic, system, sustain, integr, research, develop, provid, framework, process, review, manag, human, base, understand, approach, natur, function, impact | Seddon, Nathalie (2016) Biodiversity in the Anthropocene: prospects and policy |
| Fire & disturbance | FD | 28 | 1033 | 360 | fire, disturb, year, term, time, long, sever, recoveri, dynam, histor, increas, post, state, regim, burn, period, resili, past, follow, decad | Yao, Ying (2022) Evaluation of ecosystem resilience to drought based on drought intensity and recovery time |
| Forests & forestry | FO | 37 | 1322 | 912 | forest, tree, structur, stand, tropic, growth, regener, plantat, canopi, speci, densiti, natur, site, plot, pine, old, disturb, manag, studi, age | Osorio-Salomon, Karina (2021) Accelerating tropical cloud forest recovery: Performance of nine late-successional tree species |
| Freshwater & wetlands | FW | 56 | 1409 | 870 | water, river, wetland, lake, fish, flow, stream, freshwat, riparian, aquat, qualiti, condit, basin, hydrolog, flood, studi, pond, result, watersh, season | Miller, Kimberly A. (2013) Environmental Flows Can Reduce the Encroachment of Terrestrial Vegetation into River Channels: A Systematic Literature Review |
| Global change biology | GC | 27 | 1342 | 642 | chang, climat, temperatur, respons, futur, impact, increas, adapt, global, condit, shift, effect, scenario, warm, will, drought, vulner, extrem, sensit, project | Jones, Kendall R. (2016) Incorporating climate change into spatial conservation prioritisation: A review |
| Human-wildlife interactions | HU | 17 | 399 | 1823 | human, wildlif, predat, anim, wild, mammal, larg, prey, conflict, activ, trade, use, conserv, trap, carnivor, impact, hunt, popul, food, livestock | Wang, Sonam Wangyel (2006) Farmer attitudes towards conservation in Jigme Singye Wangchuck National Park, Bhutan |
| Invasion biology | IN | 104 | 1831 | 812 | plant, speci, invas, nativ, seed, non, alien, effect, interact, control, establish, exot, pollin, invad, impact, dispers, bee, grass, competit, success | Lach, Lori (2010) Contrasting effects of an invasive ant on a native and an invasive plant |
| Landscapes & land use | LU | 21 | 630 | 913 | land, use, cover, veget, grassland, agricultur, landscap, manag, increas, natur, graze, type, effect, intens, product, field, farm, habitat, crop, result | Maron, Martine (2007) Agricultural intensification and loss of matrix habitat over 23 years in the West Wimmera, south-eastern Australia |
| Marine & coastal ecology | ME | 42 | 1900 | 875 | marin, fish, coastal, reef, coral, sea, ocean, fisheri, mangrov, structur, food, trophic, impact, studi, web, level, estuari, coast, benthic, increas | Perry, Chris T. (2019) Changing geo-ecological functions of coral reefs in the Anthropocene |
| Protected areas | PA | 8 | 302 | 1338 | conserv, area, biodivers, protect, nation, global, identifi, region, reserv, prioriti, threat, loss, plan, countri, threaten, assess, current, hotspot, need, within | Oldekop, J. A. (2016) A global assessment of the social and conservation outcomes of protected areas |
| Regional studies | RG | 2 | 407 | 841 | region, studi, ecolog, factor, spatial, area, pattern, analysi, china, signific, show, veget, result, indic, differ, zone, relationship, index, mountain, trend | Ran, Chen (2021) Evaluation of temporal and spatial changes of global ecosystem health |
| Soil carbon & nutrients | SC | 62 | 1621 | 680 | soil, carbon, nutrient, organ, effect, biomass, mine, increas, product, concentr, signific, ecosystem, nitrogen, microbi, rate, cycl, studi, litter, properti, content | Lal, Rattan (2018) Digging deeper: A holistic perspective of factors affecting soil organic carbon sequestration in agroecosystems |
| Species distributions | SD | 4 | 423 | 941 | speci, model, habitat, distribut, use, rang, predict, suitabl, spatial, variabl, scale, occurr, nich, potenti, environment, import, select, result, data, across | Jha, Kaushalendra Kumar (2020) Habitat suitability mapping for migratory and resident vultures: A case of Indian stronghold and species distribution model |
| Threatened species | TS | 18 | 590 | 1805 | speci, popul, genet, island, extinct, conserv, threaten, endem, endang, list, red, amphibian, new, risk, record, collect, two, host, status, rang | Gomez, Jose M. (2015) The silent extinction: climate change and the potential hybridization-mediated extinction of endemic high-mountain plants |
| Urban ecology | UR | 43 | 992 | 859 | urban, landscap, habitat, connect, area, fragment, green, citi, patch, network, space, natur, studi, within, ecolog, environ, effect, biodivers, road, corridor | Jung, Kirsten (2018) Trait-dependent tolerance of bats to urbanization: a global meta-analysis |
| Wildlife ecology | WI | 8 | 430 | 1597 | popul, bird, site, individu, size, surviv, breed, rate, nest, densiti, declin, season, transloc, abund, habitat, year, reproduct, forag, bat, estim | Crosby, Andrew D. (2015) Looking beyond rare species as umbrella species: Northern Bobwhites (*Colinus virginianus*) and conservation of grassland and shrubland birds |

**Table S2:** Subtopics of the "Conservation science" topic *as identified using a Latent Dirichlet Allocation model run on the titles, keywords and abstracts of each paper. Core papers used the term "novel ecosystem", citing papers cited at least one core paper and applied ecology sample reflect count from a general paper search to capture topics that did not contain substantial NEC literature. Subtopic terms were the top 20 terms used more within a topic than across topics. Subtopic terms and example papers were used to name each subtopic.*

| **Topic name** | **Core paper count** | **Citing paper count** | **Applied ecology sample count** | **Subtopic-associated words** | **Example paper** |
| --- | --- | --- | --- | --- | --- |
| Bibliometrics | 3 | 158 | 151 | research, scienc, review, scientif, literatur, studi, polici, field, question, paper, gap, analysi, evid, scientist, challeng | Romanelli, Joao Paulo (2021) Four challenges when conducting bibliometric reviews and how to deal with them |
| Citizen science | 0 | 60 | 189 | particip, engag, project, scienc, program, citizen, educ, learn, monitor, success, communic, data, activ, environment, base | Tulloch, Ayesha I. T. (2013) Realising the full potential of citizen science monitoring programs |
| Conservation & climate change | 7 | 138 | 69 | chang, climat, adapt, sustain, global, futur, challeng, impact, develop, resili, solut, need, strategi, polici, capac | Bellwood, David R. (2019) Coral reef conservation in the Anthropocene: Confronting spatial mismatches and prioritizing functions |
| Conservation & livelihoods | 1 | 57 | 193 | communiti, forest, local, livelihood, base, studi, use, sustain, resourc, rural, sic, household, interview, benefit, reserv | Wang, Chengchao (2016) Assessment of sustainable livelihoods of different farmers in hilly red soil erosion areas of southern China |
| Conservation attitudes | 0 | 99 | 171 | percept, attitud, survey, influenc, public, factor, respond, perceiv, toward, differ, environment, valu, result, behavior, support | Hartig, Terry (2007) Psychological restoration in nature as a source of motivation for ecological behaviour |
| Conservation ethics | 11 | 138 | 100 | valu, anim, ethic, practic, differ, mean, debat, concept, argu, consid, one, concern, mani, moral, discuss | Vucetich, John A. (2015) Evaluating whether nature's intrinsic value is an axiom of or anathema to conservation |
| Conservation in the Anthropocene | 18 | 221 | 93 | human, environment, polit, relat, new, paper, justic, concept, way, anthropocen, explor, examin, world, ecolog, histori | Malhi, Yadvinder (2017) The Concept of the Anthropocene |
| Conservation planning | 3 | 65 | 153 | social, stakehold, process, use, approach, decis, plan, model, make, inform, integr, framework, assess, network, differ | Biggs, Duan (2011) The implementation crisis in conservation planning: could "mental models" help? |
| Ecosystem Services | 5 | 124 | 165 | ecosystem, land, servic, biodivers, polici, econom, privat, agricultur, farmer, environment, use, sustain, market, develop, increas | Asah, Stanley T. (2014) Perception, acquisition and use of ecosystem services: Human behavior, and ecosystem management and policy implications |
| Human-wildlife conflict | 0 | 38 | 190 | conserv, wildlif, conflict, human, support, trade, popul, potenti, reintroduct, understand, effort, hunt, import, coexist, polici | Dickman, A. J. (2010) Complexities of conflict: the importance of considering social factors for effectively resolving human-wildlife conflict |
| Indigenous perspectives | 8 | 153 | 91 | ecolog, restor, landscap, knowledg, cultur, ecosystem, indigen, rewild, practic, tradit, system, peopl, divers, social, valu | Wehi, Priscilla M. (2017) Importance of including cultural practices in ecological restoration |
| Perspectives on invasive species | 2 | 112 | 142 | speci, invas, plant, use, public, nativ, tree, bird, knowledg, non, inform, control, increas, biolog, threat | Genovart, Meritxell (2013) Holding up a mirror to the society: Children recognize exotic species much more than local ones |
| Resource management | 2 | 78 | 115 | manag, govern, resourc, institut, state, collabor, water, river, fish, agenc, scale, organ, fisheri, challeng, common | Bonnell, Joseph E. (2007) Stumbling forward: The organizational challenges of building and sustaining collaborative watershed management |
| Social value of protected areas | 0 | 51 | 145 | area, protect, nation, marin, studi, park, develop, coastal, site, wetland, import, use, tourism, region, island | Dudley, Nigel (2009) The Links between Protected Areas, Faiths, and Sacred Natural Sites |
| Urban biodiversity | 6 | 113 | 103 | natur, urban, biodivers, peopl, well, human, experi, citi, health, relationship, space, green, connect, contribut, free | Shwartz, Assaf (2014) Enhancing urban biodiversity and its influence on city-dwellers: An experiment |

**Table S3:** Subtopics of the "Ecosystem services" topic *as identified using a Latent Dirichlet Allocation model run on the titles, keywords and abstracts of each paper. Core papers used the term "novel ecosystem", citing papers cited at least one core paper and applied ecology sample reflect count from a general paper search to capture topics that did not contain substantial NEC literature. Subtopic terms were the top 20 terms used more within a topic than across topics. Subtopic terms and example papers were used to name each subtopic.*

| **Topic name** | **Core paper count** | **Citing paper count** | **Applied ecology sample count** | **Subtopic-associated words** | **Example paper** |
| --- | --- | --- | --- | --- | --- |
| Bibliometrics | 3 | 96 | 33 | research, studi, review, literatur, gap, systemat, analysi, focus, identifi, knowledg, futur, field, scientif, articl, public | Shen, Zhentao (2023) Ecological restoration research progress and prospects: A bibliometric analysis |
| Biodiversity conservation | 3 | 44 | 37 | biodivers, conserv, global, need, polici, challeng, approach, integr, will, target, loss, divers, potenti, requir, key | Isbell, Forest (2015) The biodiversity-dependent ecosystem service debt |
| Complex ecological systems | 4 | 62 | 49 | model, ecolog, scale, dynam, process, complex, network, data, spatial, integr, base, understand, across, use, predict | Lau, Matthew K. (2017) Ecological network metrics: opportunities for synthesis |
| Ecological interactions | 5 | 119 | 27 | function, communiti, interact, plant, speci, trait, effect, divers, understand, organ, level, structur, popul, influenc, import | Brooker, Rob W. (2008) Facilitation in plant communities: the past, the present, and the future |
| Ecosystem service indicators | 1 | 57 | 63 | assess, indic, use, ecosystem, health, framework, evalu, wetland, account, measur, method, condit, data, environment, approach | Mononen, L. (2016) National ecosystem service indicators: Measures of social-ecological sustainability |
| Ecosystem services research | 5 | 92 | 101 | ecosystem, servic, valu, provis, cultur, manag, regul, provid, benefit, well, support, decis, econom, use, import | Bennett, Elena M. (2009) Understanding relationships among multiple ecosystem services |
| Forests & agriculture | 5 | 77 | 52 | forest, landscap, land, use, product, manag, agricultur, food, increas, multifunct, result, tree, tropic, local, relat | Meyfroidt, Patrick (2011) Global Forest Transition: Prospects for an End to Deforestation |
| Historical ecology | 19 | 114 | 22 | ecolog, human, concept, process, term, scienc, time, new, conceptu, theori, histor, long, anthropocen, discuss, present | Ellis, Erle C. (2015) Ecology in an anthropogenic biosphere |
| Marine & coastal ecosystem services | 4 | 114 | 54 | manag, ecosystem, marin, coastal, freshwat, river, human, impact, habitat, stressor, effect, aquat, reef, activ, multipl | Giakoumi, Sylvaine (2015) Towards a framework for assessment and management of cumulative human impacts on marine food webs |
| Nature-based solutions | 8 | 112 | 15 | urban, natur, base, solut, design, citi, plan, nbs, green, benefit, infrastructur, provid, sustain, contribut, develop | Pineda-Pinto, Melissa (2022) The potential of nature-based solutions to deliver ecologically just cities: Lessons for research and urban planning from a systematic literature review |
| Novel ecosystem management | 38 | 82 | 16 | restor, ecosystem, ecolog, speci, invas, impact, novel, biolog, non, nativ, degrad, new, veget, habitat, success | Hobbs, RJ (2006) Novel ecosystems: theoretical and management aspects of the new ecological world order |
| Resilience under climate change | 10 | 110 | 41 | chang, climat, resili, ecosystem, manag, disturb, adapt, risk, global, respons, state, natur, driver, shift, increas | Weiskopf, Sarah R. (2020) Climate change effects on biodiversity, ecosystems, ecosystem services, and natural resource management in the United States |
| Social-ecological management | 3 | 85 | 50 | system, social, manag, ecolog, framework, approach, socio, govern, transform, integr, decis, complex, practic, make, adapt | Westley, Frances R. (2013) A Theory of Transformative Agency in Linked Social-Ecological Systems |
| Supply and demand in ecosystem services | 1 | 45 | 108 | area, suppli, trade, studi, ecolog, demand, off, soil, carbon, region, spatial, scenario, analysi, water, land | Sturck, Julia (2014) Mapping ecosystem services: The supply and demand of flood regulation services in Europe |
| Sustainable development | 4 | 95 | 38 | sustain, develop, environment, water, econom, resourc, energi, goal, impact, technolog, innov, achiev, integr, secur, polici | Gaspar, Jorge dos Santos (2017) The traditional energy-growth nexus: A comparison between sustainable development and economic growth approaches |

***Table S4:*** *Negative binomial model summary predicting the count of citing paper counts by the number of core Novel Ecosystem papers within each topic. Core paper count was natural-log transformed, such that the slope represents the % increase in citing paper count expected for a 1% increase in core paper count.*

| **Fixed effects** | **Observed model** | | |  |
| --- | --- | --- | --- | --- |
| Variable | Estimate | Std Error | z-value | *p*-value |
| Intercept | 5.329 | 0.211 | 25.277 | < 0.001 |
| Core paper count | 0.466 | 0.062 | 7.461 | < 0.001 |
| **Model diagnostic tests** | |  |  |  |
| Test name | Statistic | p-value |  |  |
| Uniformity | 0.169 | 0.426 |  |  |
| Dispersion | 0.832 | 0.808 |  |  |

***Table S5:*** *Model summary for topic publication growth model. Model used negative binomial distribution and interceptless formula to model separate topic intercepts and linear and quadratic growth rate slopes. The "years from 2006" fixed effect was a continuous integer ranging from 0 (2006) to 19 (2025).*

| Fixed effect | Estimate | Std Error | z-value | *p*-value |
| --- | --- | --- | --- | --- |
| Soil carbon & nutrients (SC) | -0.309 | 0.303 | -1.022 | 0.307 |
| Marine & coastal ecology (ME) | -0.688 | 0.321 | -2.142 | 0.032 |
| Ecological restoration (ER) | 0.351 | 0.274 | 1.281 | 0.200 |
| Threatened species (TS) | -0.331 | 0.397 | -0.834 | 0.404 |
| Human-wildlife interations (HU) | -3.023 | 0.786 | -3.846 | 0.000 |
| Community & functional ecology (CE) | -0.151 | 0.304 | -0.498 | 0.619 |
| Protected areas (PA) | -0.670 | 0.492 | -1.361 | 0.173 |
| Freshwater & wetlands (FW) | 0.265 | 0.264 | 1.003 | 0.316 |
| Species distributions (SD) | -0.480 | 0.426 | -1.128 | 0.259 |
| Landscapes & land use (LU) | 0.150 | 0.315 | 0.477 | 0.633 |
| Global change biology (GC) | 0.453 | 0.254 | 1.783 | 0.075 |
| Invasion biology (IN) | 1.481 | 0.180 | 8.229 | < 0.001 |
| Urban ecology (UR) | -0.180 | 0.340 | -0.531 | 0.595 |
| Ecosystem services (ES) | 0.802 | 0.236 | 3.404 | 0.001 |
| Forests & forestry (FO) | 0.215 | 0.273 | 0.785 | 0.432 |
| Wildlife ecology (WI) | -1.857 | 0.611 | -3.039 | 0.002 |
| Fire & disturbance (FD) | -0.865 | 0.368 | -2.351 | 0.019 |
| Conservation science (CS) | -0.116 | 0.288 | -0.401 | 0.688 |
| Regional studies (RG) | -2.621 | 0.806 | -3.253 | 0.001 |
| Ecological modelling &statistics (EM) | -0.567 | 0.396 | -1.430 | 0.153 |
| yearsFrom2006:SC | 0.633 | 0.051 | 12.360 | < 0.001 |
| yearsFrom2006:ME | 0.061 | 0.074 | 0.825 | 0.409 |
| yearsFrom2006:ER | -0.094 | 0.071 | -1.314 | 0.189 |
| yearsFrom2006:TS | -0.179 | 0.085 | -2.094 | 0.036 |
| yearsFrom2006:HU | 0.143 | 0.134 | 1.069 | 0.285 |
| yearsFrom2006:CE | -0.037 | 0.073 | -0.503 | 0.615 |
| yearsFrom2006:PA | -0.192 | 0.101 | -1.905 | 0.057 |
| yearsFrom2006:FW | -0.064 | 0.069 | -0.933 | 0.351 |
| yearsFrom2006:SD | -0.149 | 0.091 | -1.645 | 0.100 |
| yearsFrom2006:LU | -0.122 | 0.077 | -1.590 | 0.112 |
| yearsFrom2006:GC | -0.087 | 0.068 | -1.277 | 0.202 |
| yearsFrom2006:IN | -0.169 | 0.061 | -2.760 | 0.006 |
| yearsFrom2006:UR | -0.122 | 0.077 | -1.582 | 0.114 |
| yearsFrom2006:ES | -0.157 | 0.067 | -2.355 | 0.019 |
| yearsFrom2006:FO | -0.072 | 0.070 | -1.023 | 0.306 |
| yearsFrom2006:WI | -0.006 | 0.112 | -0.057 | 0.955 |
| yearsFrom2006:FD | 0.048 | 0.080 | 0.596 | 0.551 |
| yearsFrom2006:CS | -0.019 | 0.071 | -0.272 | 0.785 |
| yearsFrom2006:RG | 0.003 | 0.136 | 0.020 | 0.984 |
| yearsFrom2006:EM | -0.100 | 0.084 | -1.178 | 0.239 |
| yearsFrom2006^2:SC | -0.018 | 0.002 | -8.652 | < 0.001 |
| yearsFrom2006^2:ME | -0.002 | 0.003 | -0.570 | 0.569 |
| yearsFrom2006^2:ER | 0.000 | 0.003 | 0.154 | 0.878 |
| yearsFrom2006^2:TS | 0.007 | 0.003 | 2.052 | 0.040 |
| yearsFrom2006^2:HU | -0.003 | 0.005 | -0.672 | 0.502 |
| yearsFrom2006^2:CE | 0.000 | 0.003 | 0.136 | 0.892 |
| yearsFrom2006^2:PA | 0.006 | 0.004 | 1.498 | 0.134 |
| yearsFrom2006^2:FW | 0.001 | 0.003 | 0.373 | 0.709 |
| yearsFrom2006^2:SD | 0.004 | 0.004 | 1.124 | 0.261 |
| yearsFrom2006^2:LU | 0.001 | 0.003 | 0.408 | 0.683 |
| yearsFrom2006^2:GC | 0.001 | 0.003 | 0.434 | 0.665 |
| yearsFrom2006^2:IN | 0.003 | 0.003 | 1.333 | 0.183 |
| yearsFrom2006^2:UR | 0.005 | 0.003 | 1.625 | 0.104 |
| yearsFrom2006^2:ES | 0.004 | 0.003 | 1.540 | 0.124 |
| yearsFrom2006^2:FO | 0.001 | 0.003 | 0.443 | 0.658 |
| yearsFrom2006^2:WI | 0.001 | 0.004 | 0.331 | 0.741 |
| yearsFrom2006^2:FD | -0.003 | 0.003 | -0.807 | 0.420 |
| yearsFrom2006^2:CS | 0.000 | 0.003 | 0.108 | 0.914 |
| yearsFrom2006^2:RG | 0.004 | 0.005 | 0.688 | 0.491 |
| yearsFrom2006^2:EM | 0.004 | 0.003 | 1.150 | 0.250 |
| **Model diagnostic tests** | | |  |  |
| Test name | Statistic | p-value |  |  |
| Dispersion test | 0.994 | 0.968 |  |  |
| Uniformity test | 0.044 | 0.419 |  |  |

*Table S6: Model summary for the "size-correction" citation count model, modelling citation counts of topic pairs across the Novel Ecosystem Literature. Fixed effects in this model were all z-score normalized (mean = 0, standard deviation = 1). This allows non-interaction effect slopes to be directly compared on the same scale (change in response for a one standard deviation increase in the fixed effect).*

| **Fixed effects** | | |  | | | |  |  |  |  |
| --- | --- | --- | --- | --- | --- | --- | --- | --- | --- | --- |
| Variable | | | Estimate | | Std Error | | | z-value | | *p*-value |
| Intercept | | | 4.994 | | 0.126 | | | 39.703 | | < 0.001 |
| NEC publication count (citing topic) | | | 0.441 | | 0.068 | | | 6.501 | | < 0.001 |
| NEC publication count (cited topic) | | | 0.597 | | 0.112 | | | 5.348 | | < 0.001 |
| Sample literature publication count (citing topic) | | | -0.073 | | 0.068 | | | -1.080 | | 0.280 |
| Sample literature publication count (cited topic) | | | -0.141 | | 0.112 | | | -1.268 | | 0.205 |
| NEC publication interaction (cited:citing) | | | 0.086 | | 0.030 | | | 2.823 | | 0.005 |
| Sample literature publication interaction (cited:citing) | | | 0.173 | | 0.028 | | | 6.102 | | < 0.001 |
| **Random effects** |  |  |  |  |  |  |  |  |  |  |
| Group | Variance | Std Dev | |  | |  | | |  |  |
| Citing topic | 0.074 | 0.272 | |  | |  | | |  |  |
| Cited topic | 0.227 | 0.476 | |  | |  | | |  |  |
| **Model diagnostic tests** |  |  | |  | |  | | |  |  |
| Test name | Statistic | p-value | |  | |  | | |  |  |
| Uniformity | 0.759 | 0.888 | |  | |  | | |  |  |
| Dispersion | 0.047 | 0.381 | |  | |  | | |  |  |

*Table S7: Model summary for the "linguistic similarity" citation count model, modelling citation counts of topic pairs across the Novel Ecosystem Literature. Fixed effects in this model were all z-score normalized (mean = 0, standard deviation = 1). This allows non-interaction effect slopes to be directly compared on the same scale (change in response for a one standard deviation increase in the fixed effect).*

| **Fixed effects** | | |  | | | |  |  |  |  |
| --- | --- | --- | --- | --- | --- | --- | --- | --- | --- | --- |
| Variable | | | Estimate | | Std Error | | | z-value | | *p*-value |
| Intercept | | | 4.924 | | 0.097 | | | 50.503 | | < 0.001 |
| NEC publication count (citing topic) | | | 0.657 | | 0.061 | | | 10.722 | | < 0.001 |
| NEC publication count (cited topic) | | | 0.822 | | 0.084 | | | 9.809 | | < 0.001 |
| Sample literature publication count (citing topic) | | | 0.100 | | 0.061 | | | 1.647 | | 0.100 |
| Sample literature publication count (cited topic) | | | 0.035 | | 0.083 | | | 0.425 | | 0.671 |
| NEC publication interaction (cited:citing) | | | 0.004 | | 0.024 | | | 0.147 | | 0.883 |
| Sample literature publication interaction (cited:citing) | | | 0.051 | | 0.023 | | | 2.201 | | 0.028 |
| Linguistic similarity between citing & cited topics | | | 0.590 | | 0.039 | | | 15.179 | | < 0.001 |
| **Random effects** |  |  |  |  |  |  |  |  |  |  |
| Group | Variance | Std Dev | |  | |  | | |  |  |
| Citing topic | 0.059 | 0.243 | |  | |  | | |  |  |
| Cited topic | 0.122 | 0.349 | |  | |  | | |  |  |
| **Model diagnostic tests** |  |  | |  | |  | | |  |  |
| Test name | Statistic | p-value | |  | |  | | |  |  |
| Uniformity | 0.618 | 0.776 | |  | |  | | |  |  |
| Dispersion | 0.039 | 0.624 | |  | |  | | |  |  |

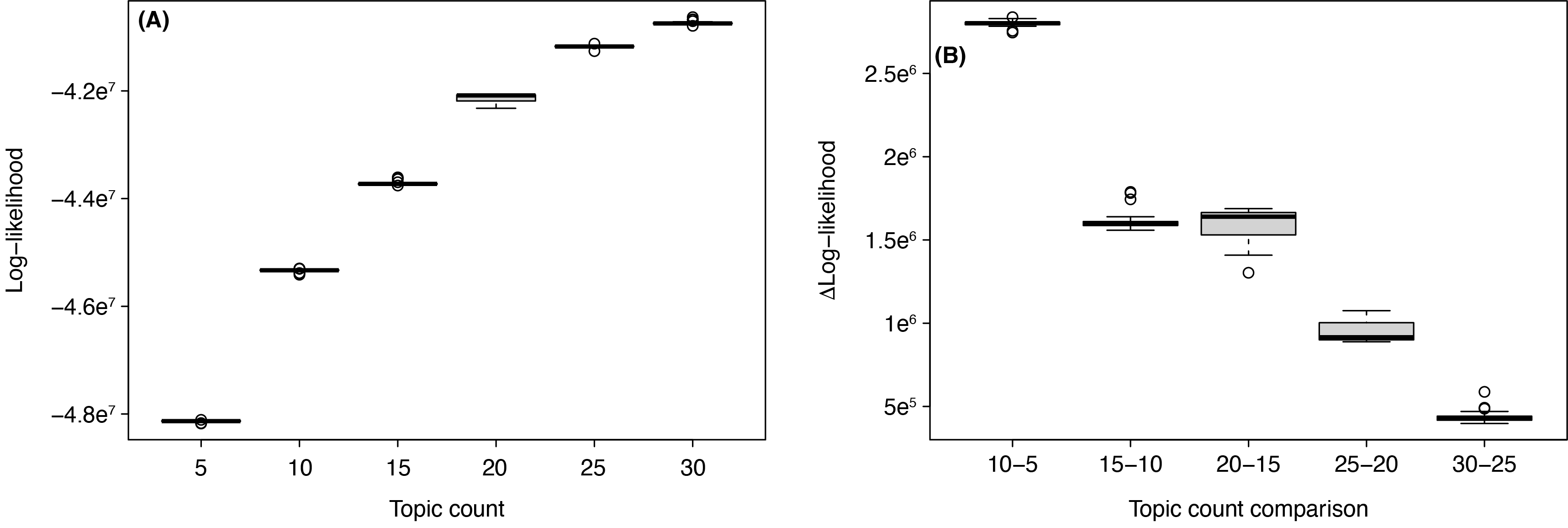

***Figure S1:*** *Validation of topic count for linguistic Latent Dirichlet Association (LDA) model used to identify topics within the novel ecosystem literature. We repeated the LDA fitted with 5-30 topics, with model performance evaluated by* ***(A)*** *log-likelihood and* ***(B)*** *pairwise change in log-likelihood. Twenty topics represented the inflection point of increases in log-likelihood, which was used in the main text models and subsequent analyses.*

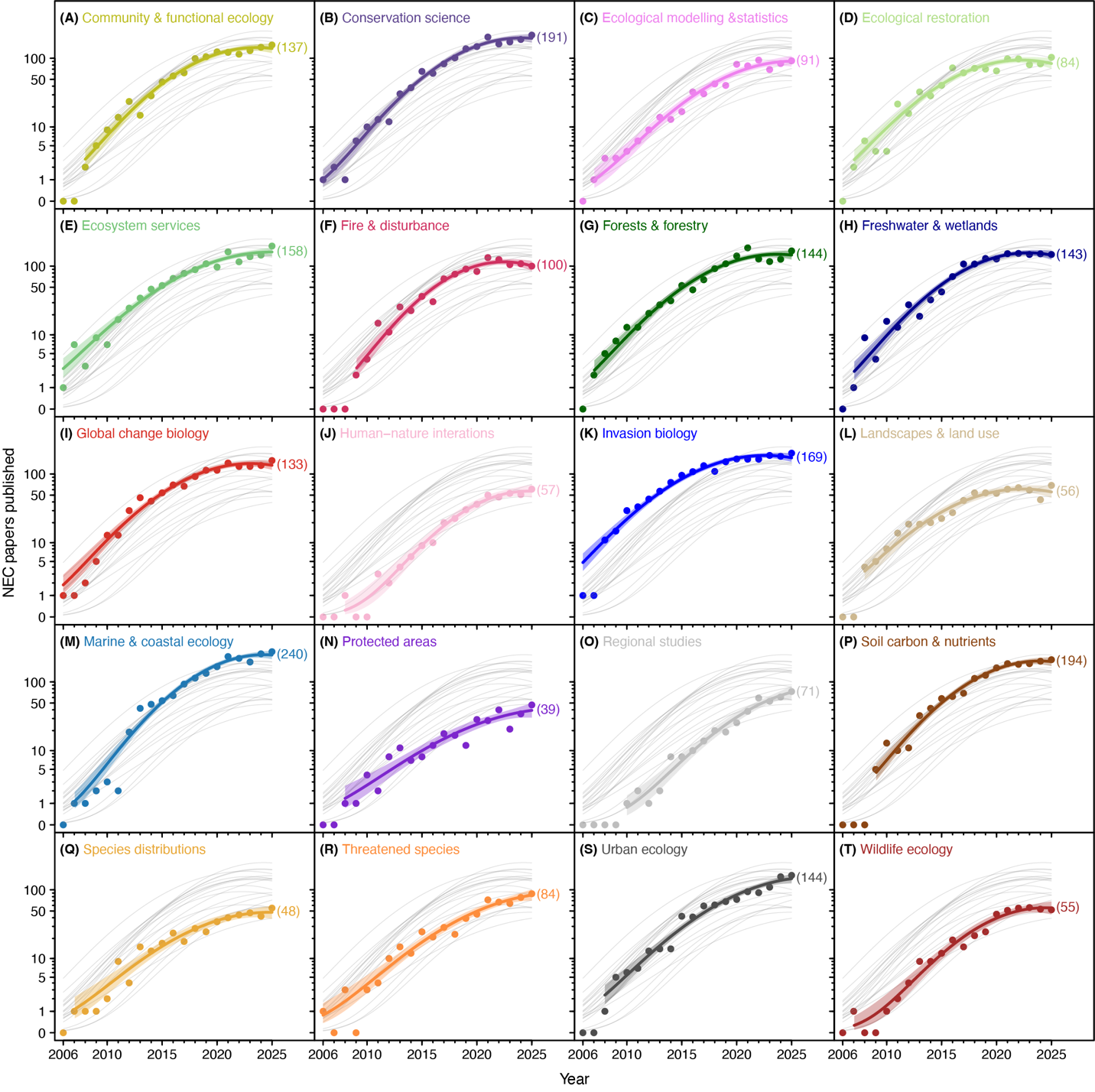
***Figure S2:*** *Growth of NEC papers (core papers using the term "novel ecosystem" and papers citing core papers) across the study period, separated into linguistic topics. Points are observed publication counts for a given year, with lines and shaded regions reflecting mean trend over time and accompanying 95% confidence intervals. Grey lines are mean trends over time for all other topics for visual comparison. Numbers in parentheses are the predicted publication count for 2025, the final year of the study period. Predicted publication count for years between 2010 and 2025 were used to establish topic rank order (Fig. 3).*

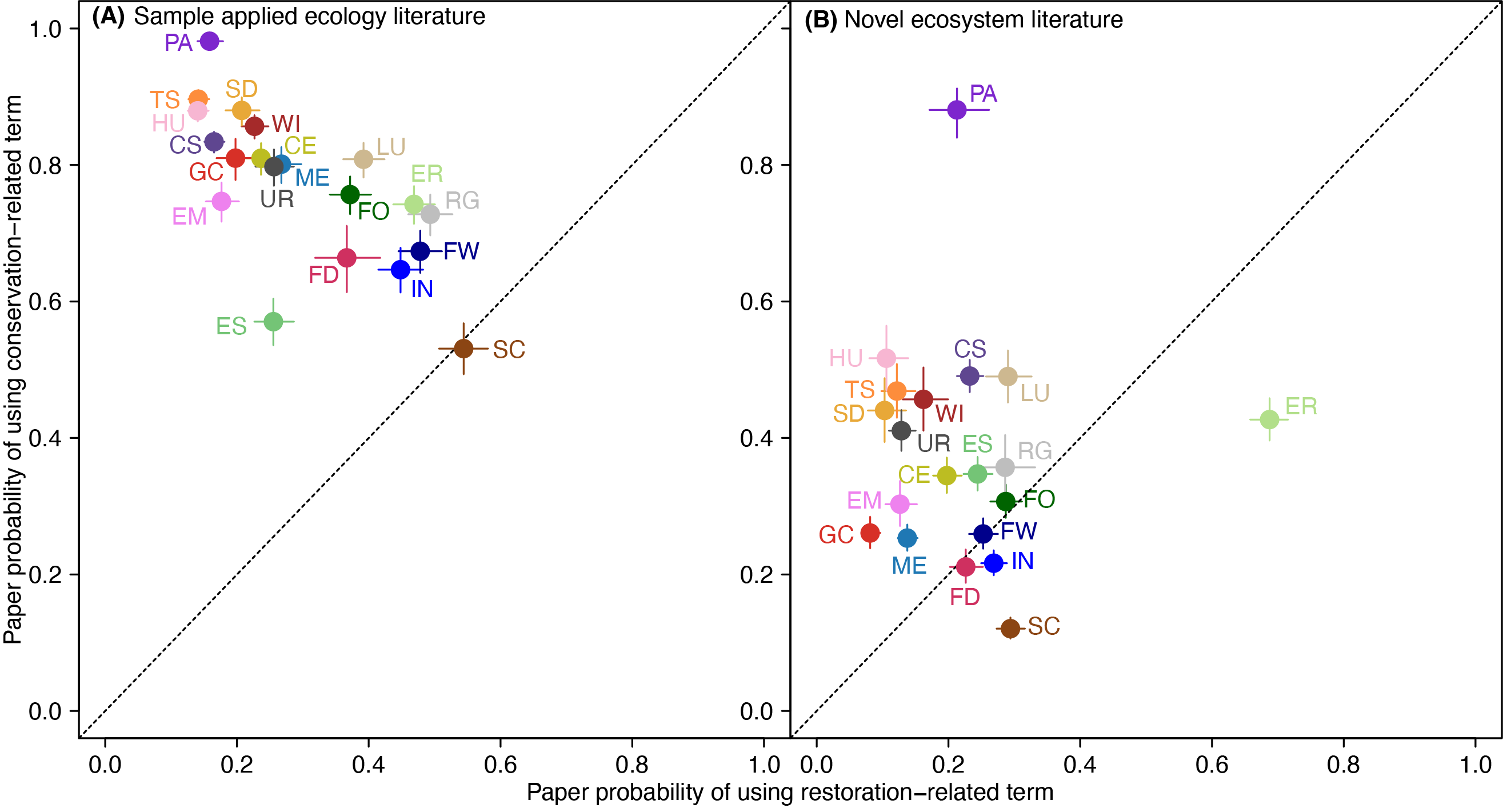

***Figure S3:*** *Probability of a paper within each content topic using a conservation-related term ("conserv", "protect" and "steward") versus a restoration-related word ("restor", "reclaim", "remediat", "reforest", "revege") within their title, keywords or abstract. (A) uses papers from the sample of applied ecology, while (B) uses papers from the NEC literature. Error bars are 95% confidence intervals. Full topic names for given acronyms are listed in Table S1.*
